# Quantitative proteomics identifies secreted diagnostic biomarkers as well as tumor-dependent prognostic targets for clear cell Renal Cell Carcinoma

**DOI:** 10.1101/2021.02.08.430238

**Authors:** Aydanur Senturk, Ayse Tugce Sahin, Ayse Armutlu, Murat Can Kiremit, Omer Acar, Selcuk Erdem, Sidar Bagbudar, Tarik Esen, Nurcan Tuncbag, Nurhan Ozlu

**Affiliations:** Department of Molecular Biology and Genetics, Koç University, 34450, Istanbul, Turkey; Department of Pathology, Koç University School of Medicine, 34010, Istanbul, Turkey; Department of Urology, Koç University School of Medicine, 34010, Istanbul, Turkey; Department of Urology, Istanbul University, Istanbul Faculty of Medicine, 34093, Istanbul, Turkey; Department of Pathology, Istanbul University, Istanbul Faculty of Medicine, 34093, Istanbul, Turkey; Graduate School of Informatics, Department of Health Informatics, Middle East Technical University, 06800, Ankara, Turkey; Cancer Systems Biology Laboratory (CanSyL), Middle East Technical University, 06800, Ankara, Turkey; Koç University Research Center for Translational Medicine (KUTTAM), 34450, Istanbul, Turkey

**Keywords:** biomarkers, drug repurposing, proteomics, Renal Cell Carcinoma, urine

## Abstract

Clear cell Renal Cell Carcinoma (ccRCC) is the third most common and most malignant urological cancer, with a 5-year survival rate of 10% for patients with advanced tumors. Here, we identified 10,160 unique proteins by in-depth quantitative proteomics, of which 955 proteins were significantly regulated between tumor and normal adjacent tissues. We verified 4 putatively secreted biomarker candidates, namely PLOD2, FERMT3, SPARC and SIRPα, as highly expressed proteins that are not affected by intra- and inter-tumor heterogeneity. Moreover, SPARC displayed a significant increase in urine samples of ccRCC patients, making it a promising marker for clinical screening assays. Furthermore, based on molecular expression profiles, we propose a biomarker panel for the robust classification of ccRCC tumors into two main clusters, which significantly differed in patient outcome with an almost three times higher risk of death for cluster 1 tumors compared to cluster 2 tumors. Moreover, among the most significant clustering proteins, 13 were targets of repurposed inhibitory FDA-approved drugs. Our rigorous proteomics approach identified promising diagnostic and tumor-discriminative biomarker candidates which can serve as therapeutic targets for the treatment of ccRCC.

## Introduction

Renal Cell Carcinoma (RCC) is among the 10 most common cancers in both men and women and causes more than 140,000 deaths worldwide every year (1). One third of the patients already display metastasis at the time of diagnosis and are expected to have a median survival of around 26 months (2, 3). Clear cell RCC (ccRCC) is the most common histological subtype accounting for more than 70% of the cases (3). Traditional treatment with chemotherapy or radiotherapy fail to treat this malignancy, limiting the treatment options primarily to surgical excision, together with targeted therapy and/or immunotherapy in suitable cases (1). Biomarkers can be used as indicators of tumor development and progression, and their discovery has largely benefitted from technological advances in high-throughput mass spectrometry-based proteomics. However, despite their urgent need, no universal biomarkers of clinical utility are currently known for ccRCC (4). This is due to previous studies lacking in coverage depth with generally less than 2,000 protein identifications and presenting insufficient validation of the candidates by multiple independent methods or in independent cohorts (5–11). Also, most candidates are restricted to the tissue context, however, biomarkers for easily available and non-invasive biospecimens such as body fluids are of more practical utility in the clinical setting (12, 13). Urine has the advantage over blood to be collectable in large amounts, to be less prone to proteolytic degradation and to be less complex, however, reliable urine biomarkers are still scarce (12, 14–16).

Over the past years, it also became clear that a “one-size-fits-all” treatment strategy for RCC patients is insufficient for the effective management of the disease and the prevention of recurrence. This is particularly due to genetic and epigenetic interpatient tumor heterogeneity caused by branched clonal evolution of the cancer cells leading to poor treatment response or to treatment resistance (4, 17). Another crucial factor causing tumor diversity is the composition of the tumor microenvironment, which consists of the extracellular matrix and different types of stromal and immune cells (18). In recent years, patient classification based on in-depth molecular expression profiles rather than histological parameters have grown in importance.

Drug treatment according to individual molecular aberrations opens new avenues for a tumor-oriented intervention. Especially in the case of grade 2 and 3 tumors, the disease progress and outcome can vary dramatically (19), and therefore be difficult to predict, which further necessitates appropriate tumor stratification. In the past years, several genomics and transcriptomics based classification systems have been proposed for ccRCC (20), such as the microarray derived ccA/ccB clustering signature, consisting of 34 marker genes (ClearCode34) (21). This clustering signature revealed noticeable prognostic difference between the low-risk ccA tumors, characterized particularly by fatty acid metabolism and angiogenesis, and the high-risk ccB tumors, associated with EMT and Wnt signaling. However, classification systems of RCC based on protein expression data have rarely been described (20, 22).

Here, we performed global quantitative shotgun proteomics with ccRCC tissues and validated potential biomarker candidates by taking orthogonal approaches. We also discovered a novel secreted biomarker in urine samples. We further developed a two-cluster based classification system for the risk stratification of ccRCC patients using the generated proteome profiles.

## Material and Methods

### Ethics statement

The study proposal was approved by the Ethics Committee of Koç University in September 2017 (no. 2017.145.IRGB2.051). The approval was prolonged for 2 more years and was expanded for urine sample collection in 2019.

### Sample collection

Tumor and NAT samples were collected during partial or radical nephrectomy with the patients’ approved consent in 2 hospitals in Istanbul (Koç University School of Medicine and Istanbul University Istanbul Faculty of Medicine). Samples were immediately stored at −80 °C after collection. The age of the patients varied between 28 and 74 and reflected a typical gender representation for RCC with 2.6:1 male-to-female-ratio. Urine samples were collected pre-operatively with the patients’ approved consent from ccRCC patients and a tumor-free control group in the Urology Department of the Koç University School of Medicine. Samples were supplemented with complete EDTA-free protease inhibitor and immediately stored at −80 °C. Detailed clinical information about the patient cohort is summarized in Supplementary File 1.

### In-solution protein digest of tissue samples

Normal adjacent and tumor tissue samples from 13 patients were subjected to protein isolation using 8 M urea, 1 mM sodium orthovanadate, complete EDTA-free protease inhibitor mixture (0.5 tablet for 5 mL, Thermo Fisher Scientific), phosSTOP phosphatase inhibitor mixture (0.5 tablet for 5 mL, Roche) and 1% n-octylglucoside in 50 mM ammonium bicarbonate (ABC, pH 8.0). For extensive protein extraction, 20-30 mg of tissue samples were homogenized with the Bullet Blender Tissue Homogenizer (Next Advance) using 0.5 mm zirconium oxide beads. The cells were further disrupted by passing the sample through a fine syringe needle (26G) 10 times on ice. The protein concentration was measured using the BCA assay (Pierce, Thermo Fisher Scientific), followed by reduction of disulfide bonds with 10 mM dithiothreitol (DTT) for 1 h at 56 °C and cysteine alkylation with 20 mM iodoacetamide (IAA) for 45 min in the dark at room temperature. The urea concentration was reduced to 1 M with 50 mM ABC and trypsin was added at 1:50 enzyme:protein ratio. The protein digest took place at 37 °C overnight and was quenched by acidification using 10% formic acid (FA). The protein samples were desalted on SepPak C18 cartridges (Waters) using 0.1% FA for washing (RP A solution) and 0.1% FA in 80% acetonitrile (ACN) for elution (RP B solution).

### Isotopic labeling and SCX fractionation of peptides

Peptide samples were labeled at their primary amines with stable dimethyl isotopes according to the protocol described by Boersema *et al*. (23) For light labeling 4% formaldehyde solution (CH^2^O) and 0.6 M cyanoborohydride solution (NaBH3CN) were mixed in 50 mM sodium phosphate buffer (pH 7.5), while for heavy labeling 4% ^13^CD_2_-labeled formaldehyde solution (^13^CD_2_O) and 0.6 M cyanoborodeuteride solution (NaBD3CN) were mixed. For 7 patients, the NAT samples were light labeled while the tumor tissue samples were heavy labeled. For the remaining 6 patients, the labeling was swapped in order to eliminate bias from the label choice. A total of 600-700 μg of peptides were loaded on 1cc SepPak C18 cartridges and flushed with the prepared light and heavy labeling solution, respectively. The labeled peptides were washed twice with RP A solution and eluted with RP B solution. The peptide mixtures were then fractionated on-column by Strong Cation Exchange (SCX) chromatography. The dried peptide samples were reconstituted in SCX loading buffer (7 mM KH2PO4, 30% ACN, pH 2.65) and light and heavy labeled peptides were mixed at 1:1 ratio based on their median peptide intensities. The SCX resin packed cartridge (HyperSep, Thermo Fisher Scientific) was prepared by sequential addition of following solutions: methanol, elution buffer 9, HPLC-grade water, equilibration buffer (50 mM K2HPO4, 500 mM NaCl, pH 7.5), HPLC-grade water and loading buffer. The bound peptides were eluted by step-wise addition of elution buffers which consisted of the loading buffer with increasing KCl concentrations (F1: 30 mM; F2: 45 mM; F3: 60 mM; F4: 75 mM; F5: 90 mM; F6: 105 mM; F7: 120 mM; F8: 150 mM and F9: 500 mM). The obtained 10 fractions (flow-through included) were extensively desalted using 1cc SepPak columns.

### Data acquisition

The collected SCX fractions were reconstituted in 5% FA and 5% ACN (MS analysis solution) and analyzed in duplicate with 120 min linear gradients on an UltiMate 3000 RSLCnano reversed phase chromatographic platform (Thermo Fisher Scientific) coupled to a Q Exactive hybrid quadrupole-Orbitrap mass spectrometer (Thermo Fisher Scientific). Approximately 300 ng of each fraction was loaded onto an in-house packed 100 μm i.d. × 17 cm C18 column (Reprosil-Gold C18, 5 μm, 200Å, Dr. Maisch) and run with a flow rate of 300 nL/min. The chromatographic separation of the peptides started at 4% of solution B (0.1% FA in ACN) and gradually increased to 25% in 69 min. The gradient continued from 25% to 40% of solution B in the next 20 min. Peptides in the mass range 400-1,500 m/z and with a positive polarity were allowed for detection in data-dependent acquisition mode (DDA). For the MS1 spectra acquisition the resolution was set to 70,000, the automatic gain control (AGC) target to 1e6 and the maximum injection time to 32 ms. The top 15 most intense peptides per cycle were selected for fragmentation in the higher-energy collisional dissociation (HCD) cell with a normalized collision energy (NCE) of 26. MS2 spectra acquisition was conducted at a resolution of 17,500, an AGC target of 1e6, a maximum injection time of 85 ms and a fixed first mass of 120 m/z. Furthermore, the isolation window was set to 2.0 m/z, the dynamic exclusion was set to 35 s and the charge exclusion was set as unassigned, 1, 6-8, >8.

For the targeted proteomics approach, unlabeled digest samples of patients were analyzed in Parallel Reaction Monitoring (PRM) mode. Samples were reconstituted in MS analysis solution and run with a 90 min linear gradient on a similar system as described above except that a Q Exactive HF mass spectrometer was engaged. Approximately 600 ng were loaded onto an in-house packed 100 μm i.d. × 25 cm C18 column (Reprosil-Gold C18, 3 μm, 200Å, Dr. Maisch) and run with a flow rate of 300 nL/min. The gradient started at 4% of solution B, increased to 25% in 49 min and continued from 25% to 45% of solution B in the next 15 min. Peptides with a positive polarity were targeted. MS2 spectra acquisition was performed at a resolution of 45,000, an AGC target of 2e5, a maximum injection time of 100 ms and a fixed first mass of 100 m/z. Moreover, the isolation window was set to 1.2 m/z and the NCE to 28.

### Data processing

Peptide identification and quantification was done with Proteome Discoverer (PD) (v1.4, Thermo Fisher Scientific) using the SequestHT search engine. Peptide spectral matches were searched against a Swissprot database containing 21,039 entries for *Homo sapiens* retrieved from Uniprot in March 2016. Trypsin was selected as hydrolytic enzyme with a maximum number of allowed missed cleavages of 2. Regarding peptide identification, a mass tolerance of ±20 ppm for precursor masses and ±0.05 Da for fragment ions were selected. For quantitation, the 2plex dimethyl heavy/light method with a mass precision of 2 ppm for precursor measurements was used. Light and heavy dimethylation of peptide N termini and of lysine residues, as well as methionine oxidation, were set as dynamic modifications. Cysteine carbamidomethylation was set as fixed modification. The False Discovery Rates (FDR) for peptide and protein identifications were set to 1%. Only peptides with medium or high identification confidence, with a sequence length between 7 and 25 and with a peptide rank of minimum 1 were allowed.

Quantification ratios of samples with labeling swap were transformed to heavy/light (H/L) format. H/L ratios for biological replicates were average values from two technical replicates. The obtained data was filtered for proteins quantified at least in one of the 13 patients (5,799 proteins), which are hereafter denoted as “quantified proteins”. The H/L values were sample-wise normalized by the sample median and log2 transformed.

### Urine sample preparation

Urine samples were first adjusted to pH 8.0 using 500 mM ABC supplemented with EDTA-free protease inhibitor and then filtered through an Amicon Ultra-0.5 mL Centrifugal Filter Device (10 kDa, Merck Millipore). Collection of the retentate was performed by flipping the filter device upside-down and conducting centrifugation at 1,000*g* for 2 min. The protein concentration was determined by BCA assay.

### Immunoblotting

For immunoblot analysis 40 mg of normal tissue and 70-80 mg of tumor tissue were lysed in 0.1% Triton-X in 1xPBS and EDTA-free protease inhibitor. In total, 40 μg of tissue samples and 20 μg of urine samples were run on 8-10% SDS gels and transferred to nitrocellulose membranes for 3-4 h at RT. Primary antibody incubation was done overnight at 4 °C with following dilutions: anti-PLOD2 mouse at 1:500-1:1,000 (MAB4445, R&D Systems), anti-FERMT3 mouse at 1:500-1:1,500 (ab173416, Abcam), anti-SPARC mouse at 1:1,000 (33-5500, Invitrogen), anti-SIRPα rabbit at 1:1,500-1:3,500 (13379, Cell Signaling Technology) and anti-ACTB mouse at 1:5,000 (ab6276, Abcam). As secondary antibodies HRP-conjugated anti-mouse IgG at 1:2,000 (7076S, Cell Signaling Technologies) and HRP-conjugated anti-rabbit IgG at 1:2,000 (7074S, Cell Signaling Technologies) were used. Protein expression was visualized using Clarity Western ECL Substrate (Bio-Rad). Densitometric analysis was performed with ImageJ (v1.46r) and visualization and statistical analysis (Student’s *t*-test p-value <0.05) with GraphPad PRISM v8.

### Parallel Reaction Monitoring

Selected unlabeled peptides from digest experiments were targeted by Parallel Reaction Monitoring as quantifiable surrogates for the proteins of interest. Candidate peptides were selected as follows: uniqueness of peptides was ensured by BLASTp analysis, peptides with unstable charge state across samples were dismissed, peptides with length between 7-18aa were preferred, and sequences with ragged ends, with missed cleavages and/or with post-translational modifications were avoided. When no alternative peptides were available for a protein, exceptions to these exclusion criteria were made. No exception was made for the charge state criteria. MS/MS spectra were inspected extensively and peaks with a mass window <10 ppm were integrated by manually setting the borders in Skyline (v19.1). The spectral library consisted of 60,833 spectra from DDA-based data acquisition of tumor and normal digest samples of the discovery cohort. Doubly and triply charged peptides in the range of 375-1500 m/z with a library ion match tolerance of 0.5 m/z were accepted. Only spectra with a high correlation with the spectral library (dotp value >0.8) were used. For each peptide, the summed peak areas of its 6 fragment ions was calculated and then normalized by the summed peak areas of the β-actin peptide GYSFTTTAER in the sample. Calculated tumor/normal peak area ratios were transformed to log2.

### Immunohistochemistry

Immunohistochemical analyses were performed with the BenchMark Ultra autostaining platform (Ventana Medical Systems). Briefly, the tissue paraffin sections were cut to 3 μm thickness onto charged slides. Sections were deparaffinized and rehydrated through an alcohol series. Tissue sections were incubated with primary antibodies at 37 °C with following dilutions: anti-SPARC mouse at 1:800 (33-5500, Invitrogen), anti-SIRPα rabbit at 1:25 (13379, Cell Signaling Technology), anti-FERMT3 mouse at 1:2,500 (ab173416, Abcam) and anti-PLOD2 mouse at 1:250 (MAB4445, R&D Systems). The UltraView Detection kit (760-500, Ventana Medical Systems) was used for detection. The reaction product was visualized with 3, 3′-diaminobenzidine (DAB) chromogen and counterstained with hematoxylin. All stained slides were evaluated in a blinded fashion by a single pathologist. The intensity was scored as mild (score 1: any positivity that could be seen at high magnification), moderate (score 2: any staining in-between score 1 and 3), and marked (score 3: any staining that could be seen at low magnification). The prevalence of the staining was evaluated as score 0 (<5%), score 1 (5%-25%), score 2 (25%-75%) and score 3 (>75%).

### Statistical analysis

Significantly dysregulated proteins between tumor and normal tissues were determined using the one-sample Wilcoxon signed-rank test. For this purpose, the Python function scipy.stats.wilcoxon (v0.14.0) was extended by the feature to omit missing quantification values and was then applied gene-wise on the data. Benjamini-Hochberg adjusted *p*-values <0.05 were considered statistically significant. For the determination of statistically different cluster-discriminative proteins, two-sample Student’s *t*-test was applied. *P*-values were calculated gene-wise using the ttest_ind function of the scipy.stats module by omitting missing values (nan_policy=“omit”). *P*-values <0.05 were considered statistically significant.

### Functional annotation

Significantly regulated proteins were annotated by their role in cancer using the COSMIC Cancer Gene Census database (v91). For Gene Ontology (GO) cellular component and for pathway annotations, over-representation analysis (ORA) was applied using the WebGestalt platform (v2019) (24) with a BH-adjusted FDR of 0.02 and 0.05, respectively. Pathway annotations were combined analysis of KEGG, Reactome, Hallmark50, and Wikipathway_cancer databases. For GO biological process annotation of the significantly regulated proteins the PANTHER (v14) platform (25) was used with an FDR setting of 0.05.

### Network-based interpretation of proteomic data

Optimal subnetworks, which represent the list of significant proteins best, were reconstructed by integrating a reference human interactome with our proteomic data using the Forest module of Omics Integrator (v0.3.1) (26) and the visualization tool Cytoscape (v3.8.0) (27). The iRefWeb_2013 interactome file was used as global network reference. Nodes were annotated by GO cellular compartment using QuickGO (28), which was limited to Uniprot assignments. For multiple allocations priority was given as follows: “extracellular”, “membrane”, “nucleus”, “cytoplasm”, “other”, “unknown”. Additionally, networks were annotated with GO biological process or Reactome pathways, respectively, using the Cytoscape plug-in ClueGO (v2.5.7) (29). For cluster-discriminative proteins the network was complemented with associated inhibitory FDA-approved drugs retrieved from the Drugbank database (v5.1.7, access August 2020) (30). Chemical elements were excluded as drugs.

### Hierarchical clustering and Principal Component Analysis

Unsupervised hierarchical clustering was applied sample-wise on the global proteome data. A distance matrix was prepared by calculating the arithmetic mean of the squared Euclidean distances between samples with an in-house generated Python script. The subsequent clustering was done using the linkage function of the scipy.hierarchy module with the weight method “ward” and the distance metric “Euclidean”. Besides all quantified proteins, the high-variant proteins were also clustered, which were determined using the median_absolute_deviation function of the astropy.stats module. The calculated median absolute deviation values were further corrected by the consistency factor 1.4826 and a cut-off of 1.5 was applied. Principal Component Analysis (PCA) was employed for the first two components using the PCA function of the sklearn.decomposition module.

### Comparison with ccRCC expression data

FPKM normalized RNASeq data of The Cancer Genome Atlas KIRC cohort consisting of 539 tumor and 72 normal tissue samples was retrieved from the TCGAPortal (http://tcgaportal.org/download.html, April 2020). Genes with more than 70% missing expression values were filtered out and tumor counts were gene-wise normalized to the upper quartile of respective normal counts and log2 transformed. The CPTAC ccRCC proteome data was retrieved from the CPTAC portal (https://cptac-data-portal.georgetown.edu, November 2019). Quality control samples and the reported contaminated sample (“C3N-00314”) were excluded. Log2 tumor/normal ratios were calculated for the 80 tumors with adjacent normal sample by eliminating the relation to the TMT reference sample. Pearson and Spearman correlations were calculated using the respective Python scipy.stats functions.

### Survival analysis

For evaluation of the prognostic impact of the tumor clusters, the follow-up data of the TCGA-KIRC cohort for 534 patients was used (Firehouse Legacy, downloaded from cBioPortal, access August 2020). Survival analysis was performed using the KaplanMeierFitter and multivariate_logrank_test functions of the Python library lifelines (v0.25.0). To determine the impact of clinicopathological factors on survival, univariate and multivariate Cox regression analyses were performed using the CoxPHFitter function of the library. Statistical significance of the frequency of clinicopathological factors in the clusters was determined using the fisher_exact function of the scipy.stats library.

### Clustering validation and marker gene identification

To verify the observed clustering signature, two independent expression datasets of ccRCC (TCGA and CPTAC) were subjected to two-sided Student’s *t*-test after clustering (*p*-value <0.05). Cluster assignment was then conducted based on highest overlap between the significantly different proteins of the independent dataset clusters compared with those of the proteome-based clusters of this study. To identify the key discriminatory elements of the clustering signature (“marker genes”), a cut-off value of ±2 for the *t*-test difference and a cut-off of >2 for the -log_10_ *p*-value were set.

## Results

### Shotgun proteomics approach achieves deep coverage of ccRCC proteome

In this study, a comprehensive quantitative proteome analysis of ccRCC has been conducted to address the scarcity of reliable biomarker panels and to uncover underlying molecular phenotypes. To this end, dimethylation-based quantitative proteomics analysis with extensive homogenization of biospecimens has been performed with 13 ccRCC patient’s frozen tumor and normal adjacent tissue (NAT) samples (Figure 1A). The discovery cohort included patients of different age, grade, stage and gender (Supplementary File 1). We identified a total of 10,160 unique proteins from all samples with a False Discovery Rate (FDR) of 0.01. Protein identification and quantification were robust across patients with a coefficient of variation of approximately 6.7% for the identifications and 8.5% for the quantifications, respectively (Figure 1B). On average, 4,453 proteins were identified and 3,118 proteins were quantified in the paired samples, respectively. Abundance levels of the ccRCC proteome encompassed almost 6 orders of magnitude in dynamic range and correlated with diverse cellular compartments (Figure 1C). The most abundant proteins were associated with angiogenesis such as blood coagulation and platelet degranulation, and with translational processes such as chaperone activity and protein maturation in the ER. Notably, the low-abundance range was characterized by proteins predominantly localized to the mitochondrial matrix and mitochondrial inner membrane. In general, the distribution of tumor/normal fold change values followed a near normal pattern over more than 14 orders of magnitude with a noticeable skewness towards normal tissue regulation (Figure 1D,E).

**Figure 1.**
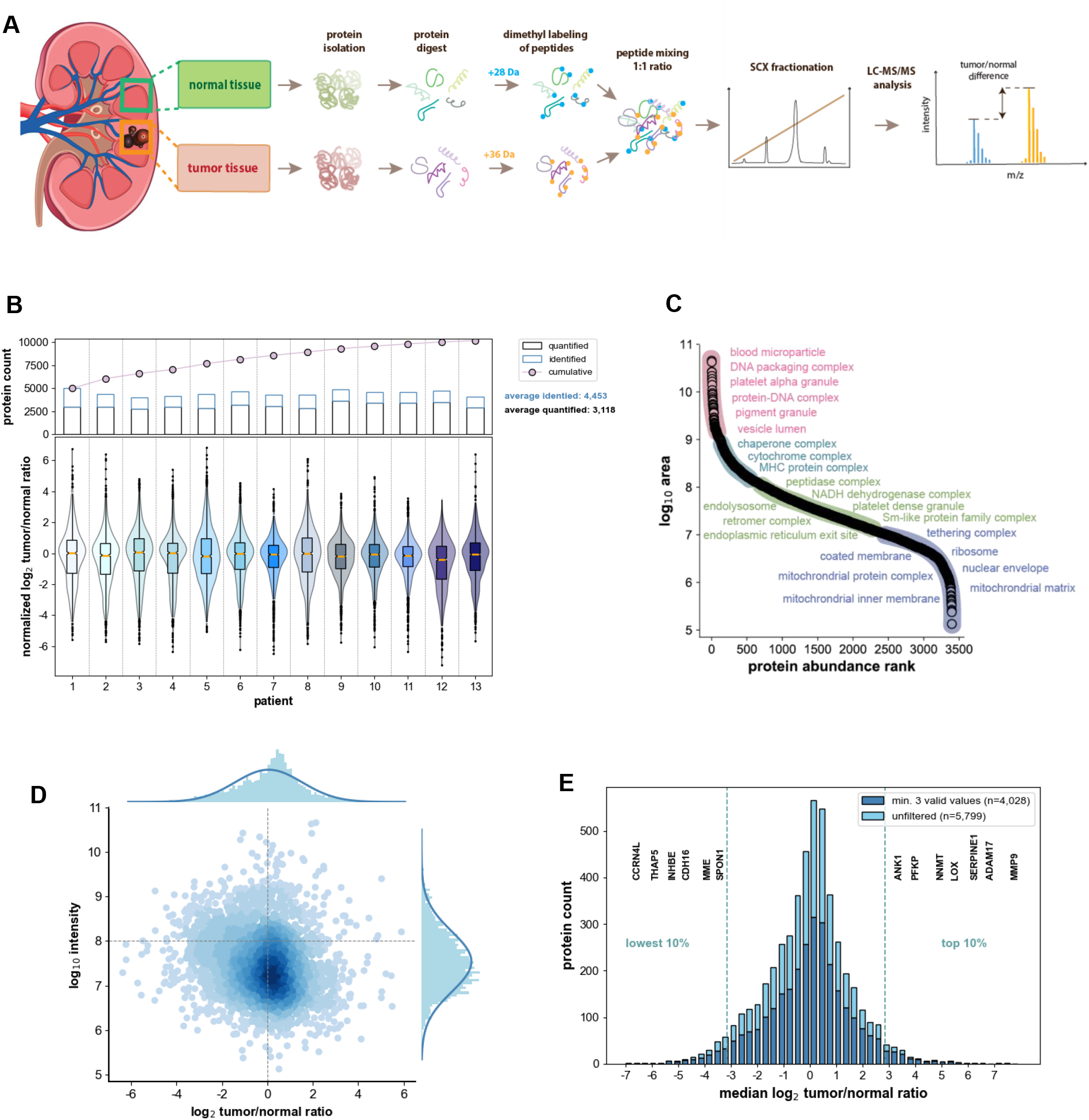
Global proteome analysis of clear cell Renal Cell Carcinoma. **(A)** Experimental outline of shotgun proteomics approach. **(B)** Number and distribution of identified and quantified proteins across all patients. **(C)** S curve showing associated cellular compartments of identified proteins. **(D)** MA plot illustrating intensity distribution of quantified proteins. **(E)** Histogram showing general distribution of median tumor/normal ratios of filtered and unfiltered quantified proteins.

### Significantly regulated proteins mirror RCC-typical metabolic alterations and molecular characteristics

In total, 5,799 of the 10,160 identified proteins were quantified by dimethyl labeling (Figure 2A). We determined 955 proteins as statistically differentially expressed between tumors and NATs (BH-adjusted *p*-value <0.05) with 409 significantly up-regulated and 546 significantly down-regulated proteins. Among these were 44 genes associated with mutations that are causally implicated in cancer, such as oncogenes, tumor suppressor genes and fusion genes, according to the COSMIC Cancer Gene Census database (Figure S1A,B). Significantly elevated oncogenes were EGFR, H3F3A, AKT1, MAPK, SRC, CALR and XPO1 (Figure 2B, Supplementary File 2). The activation of the EGFR-MAPK1 axis has previously been reported to enhance cell proliferation in RCC (22, 31). Also, our data implicated that the EGFR-PI3K-AKT1 and AKT1 related mTOR signaling pathways might be activated in ccRCC, which are both associated with tumor growth and survival (32). Another significantly up-regulated oncogene is the tyrosine kinase SRC, which has previously been reported to be recruited by EGFR and to affect its downstream players RAS and AKT1 in RCC (33). Despite its categorization as a tumor suppressor, fatty acid synthase (FAS) was highly up-regulated in the tumor tissues relative to NATs, which has previously been associated with tumor aggressiveness and poor patient outcome in RCC (34). We inferred that several oncogenes possibly mediate tumor proliferation and survival in ccRCC tumors, emphasizing the need for further investigation especially of the EGFR related molecular mechanisms in respect of alternative directed therapeutic intervention.

**Figure 2.**
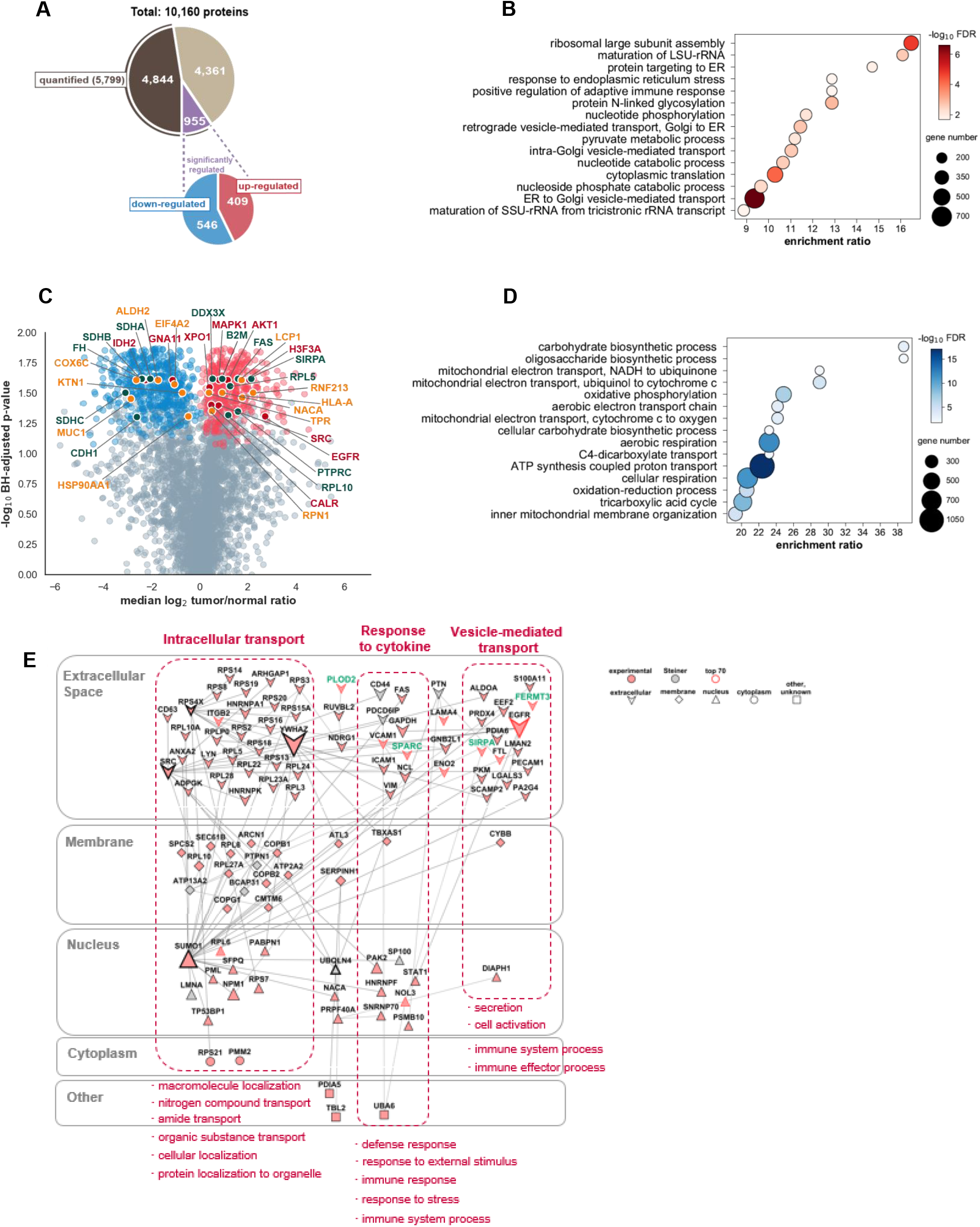
Significantly dysregulated proteins in ccRCC tumors compared to adjacent normal tissues. **(A)** Overview of the number of identified and quantified proteins. Inner pie chart represents number of significantly regulated proteins. **(B)** Volcano plot showing distribution of significantly regulated proteins with indication of gene functionality in cancer. **(C-D)** PANTHER GO-Slim Biological Process analyses of significantly up-regulated **(C)** and significantly down-regulated **(D)** proteins, respectively. **(E)** Protein-protein interaction network of significantly up-regulated proteins. Edge annotations were created using Omics Integrator by integrating intermediate interactors (Steiner nodes). Associated cellular compartment and biological process information of proteins were retrieved using QuickGO and ClueGO, respectively. Size and border width of nodes represent BetweennessCentrality. Proteins selected as biomarker candidates are indicated with green label. Network is section of most significant pathways.

As many cancers, RCC undergoes substantial metabolic reprogramming by increasing glycolytic processes and diminishing mitochondrial ATP production to meet its excessive energy requirements for tumor growth and progression (34, 35). Our findings for the significantly altered biological processes were in agreement with the reported metabolic pathobiology of RCC. The significantly up-regulated proteins were primarily associated with protein maturation and transport, PTM events such as glycosylation and phosphorylation of macromolecules, glycolysis-related processes and immune response (Figure 2C). In contrast, the significantly down-regulated proteins were mainly involved in mitochondrial events such as oxidative phosphorylation, electron transport chain, proton transport and TCA cycle (Figure 2D).

In order to unveil the most prominent cross-talks between significantly up-regulated proteins, we constructed a protein-protein interaction subnetwork using Omics Integrator (26), which confirmed that the highly elevated oncogenes EGFR and SRC were central players in ccRCC tumorigenesis (Figure 2E, Figure S2). While most of the hubs were associated with intracellular transport of macromolecules (SRC, RPS4X, SUMO1 and YWHAZ) and organelle organization (UBQLN4, Figure S2), EGFR was associated with immune system related processes and molecule secretion (Figure 2E). SUMOylation modifications have previously been reported to be elevated in a subgroup of low-grade ccRCC tumors, which were particularly associated with angiogenesis and absence of infiltrating immune cells (22). Furthermore, the regulatory protein YWHAZ (14-3-3ζ) has previously been noted to play a promoting role in primary as well as metastatic ccRCC (7, 8, 11).

### Biomarker candidates PLOD2, FERMT3, SPARC and SIRPα are highly expressed in ccRCC tumors

Given that secretion of macromolecules is essential for sculpting the tumor microenvironment to promote tumor growth and metastasis, and that vesicle-mediated transport and secretion were one of the significantly enriched biological processes in the ccRCC tumors (Figure 2), we sought to identify and characterize novel secreted biomarkers. To this end, we selected 4 biomarkers among the top 70 highly up-regulated proteins predicted to be located in the “extracellular space”. To the best of our knowledge, the selected biomarkers, namely PLOD2, FERMT3, SPARC and SIRPα, have previously not been suggested or validated as ccRCC biomarkers in proteomics-based studies. According to the CGC annotation, SIRPα has a tumor suppressor role in cancer (Figure 2B), however, in this study, SIRPα was, with a median log2 tumor/normal ratio of 2.15, clearly an up-regulated protein in the ccRCC tumor tissues.

Immunoblot analysis confirmed the elevated expression of all 4 biomarkers in the tumors compared to the adjacent normal tissues (Figure 3A). This was observed in grade 2, grade 3, as well as grade 4 patients. Next, we attempted to validate the biomarkers by targeted proteomics in digested unlabeled tissue samples. Parallel reaction monitoring allowed tracking of selected peptides, which served as surrogates for the proteins of interest, and whose product ion peak areas were used to calculate tumor/normal ratios (Figure 3B). Besides the 4 selected biomarkers, 6 further putatively secreted proteins of interest were monitored via PRM (Figure 3C,D). As expected, the selected 20 targets were more abundant in tumor samples compared to NATs and had comparable tumor/normal ratios across grades. We observed a significant positive Pearson correlation of 0.84 (*p*-value <0.001, Figure S1C) between the discovery cohort (Figure 3C) and an independent cohort consisting of 11 additional ccRCC patients (Figure 3D), which supported the suitability of these proteins as potential biomarkers.

**Figure 3.**
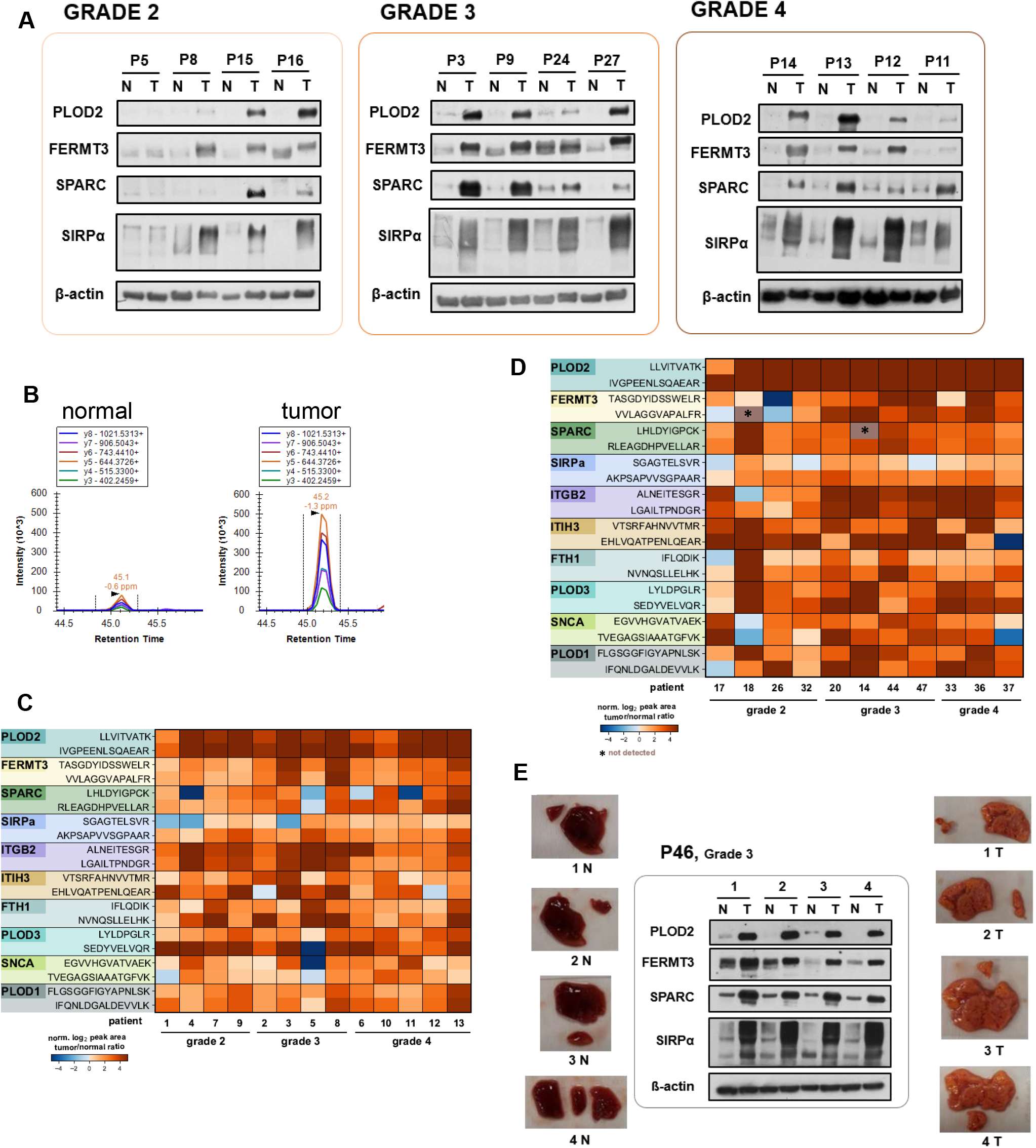
Validation of selected putative biomarkers for ccRCC. **(A)** Immunoblot analysis of biomarkers in tissues of patients with different grades. N: normal adjacent tissue, T: tumor tissue. **(B)** Targeted proteomics analysis of tissues using Parallel Reaction Monitoring. Exemplary images illustrate peak area difference of same peptide in tumor and normal adjacent sample (peptide SEDYVELVQR of PLOD3 biomarker). **(C)** Heatmap showing log2 tumor/normal ratios in PRM experiments of patient samples from discovery cohort. **(D)** Heatmap showing log2 tumor/normal ratios in PRM experiments of patient samples from independent validation cohort. **(E)** Comparable expression of biomarkers in random sub regions of tissue samples from a single patient showing independence of biomarker levels from intra-tumor heterogeneity.

Next, we tested the impact of intratumoral heterogeneity on the expression of the biomarkers in random parts of the tumor and adjacent normal tissue from a selected grade 3 patient (Figure 3E). We could not observe any remarkable local distortion of abundance for the biomarkers as evidenced by a comparable expression detected in the random sections, which further qualifies them as potentially universal biomarkers.

### SPARC localizes to the intratumoral stroma and is significantly elevated in urine of ccRCC patients

We next investigated the expression of the 4 biomarkers by immunohistochemistry (Figure 4A). Although transitional regions illustrated that the expression of PLOD2 was more intense in tumor sections (Figure S1D), a similar prevalence (score 3, >75%) was detected in both tumor and NATs (Figure 4A). SIRPα, FERMT3 and SPARC had a low prevalence score of 1 (5-25% spreading) and a low intensity score of 1 in the normal tissues, while the expression was markedly elevated and largely distributed in the tumor tissues with a prevalence score between 2 (25-75%) and 3 (>75%), and an intensity score of 3. Furthermore, for SIRPα, we determined a membranous localization in the tumor cells. Interestingly, for FERMT3 we observed almost no staining in large areas of the tumor tissues (Figure S1E), but high prevalence and intense staining in inflammatory regions (Figure 4A), coinciding with the previously reported role of FERMT3 in leukocyte transmigration, platelet degranulation and inflammation (36, 37). We were intrigued by an intense staining (score 3) for SPARC in the intratumoral stroma, which suggested that SPARC might be secreted to urine of ccRCC patients.

**Figure 4.**
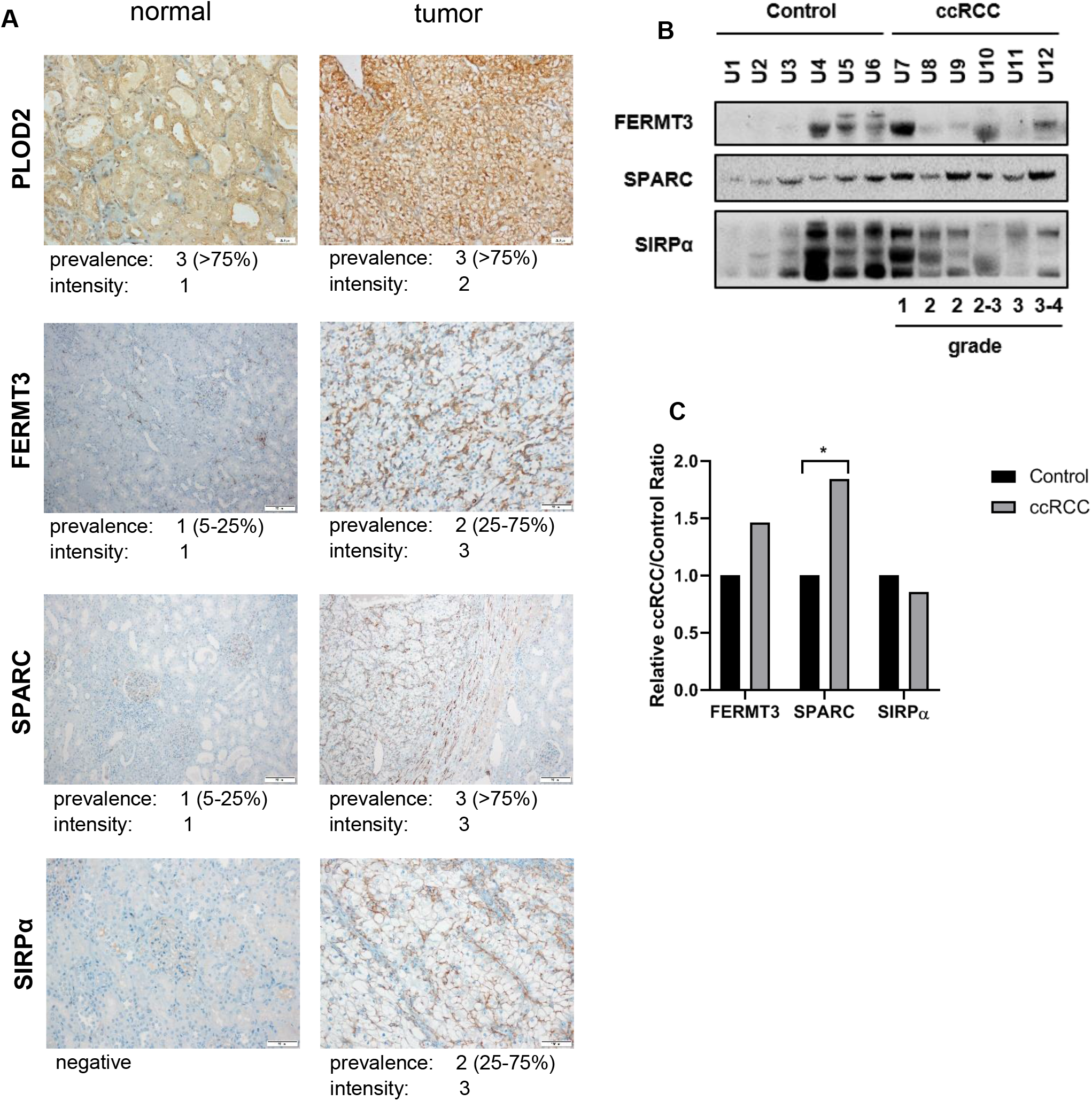
Expression profiles of selected biomarker candidates. **(A)** Immunohistochemistry staining of biomarker expression in normal and tumor tissues with prevalence and intensity scores. For SPARC, tumor picture shows transitional region to adjacent normal tissue. **(B)** Expression of biomarkers in urine samples. Control samples are from patients without tumors. **(C)** Quantification of biomarker expression in urine samples.

To challenge the suitability of the biomarkers for clinical assays, we assessed their secretion into urine, which is a readily available biospecimen and can be collected non-invasively (Figure 4B,C). We detected 3 of the 4 biomarkers in the urine samples of ccRCC patients of different grades, as well as the non-tumoral control group. Although, FERMT3 and SIRPα did not display a significant expression difference, SPARC was significantly more abundant in urine samples collected from the tumor patients in comparison to the control group (*p*-value <0.05) (Figure 4C), which makes it a promising candidate for future clinical assays.

### Molecular expression profiling stratifies ccRCC tumors into two clusters with distinct pathobiology

Our comprehensive proteomic analysis and rigorous comparison between normal and tumor tissues provided us with new biomarkers for ccRCC. Next, we focused on the inter-patient comparison in an attempt to identify a novel biomarker panel for the classification of patients based on their individual molecular expression profiles. To this end, we performed unsupervised hierarchical clustering to unveil tumor clusters. Indeed, the tumors grouped into two distinct clusters based on the expression profile of the 5,799 quantified proteins (Figure 5A). Further, we challenged the robustness of the clustering by using the expression data of 866 high-variant proteins (median absolute deviation >1.5), which recapitulated the same classification of the patients (Figure S3A). Next, we employed Principal Component Analysis (PCA), which showed concordance with the results of the hierarchical clustering and further confirmed the two main clusters (Figure 5B). Statistical analysis revealed that a total of 599 proteins were differentially expressed between clusters (Student’s *t-*test *p-*value <0.05), with 354 significantly higher abundant in cluster 1 (“cluster 1-specific”), and 245 significantly higher abundant in cluster 2 (“cluster 2-specific”), respectively (Figure 5C). Cluster 1-specific proteins were primarily involved in processes associated with the immune system, such as regulation of complement cascade and innate immune response (Figure 5D), indicating that the observed enhanced immune cell infiltration in RCC (Figure 2B,D) was primarily relevant in this subset of patients. Furthermore, platelet activation and hemostasis implied neovascularization, and therefore a concomitant angiogenic profile of these tumors. Also noteworthy is the enrichment of components associated with the regulation of the insulin-like growth factor IGF, which implied a primary dependence on this growth factor for tumor progression. In contrast, cluster 2 tumors displayed an enrichment of metabolic processes related to lipid, fatty acid and amino acid degradation (Figure 5E). Moreover, proteins associated with mitochondrial processes such as oxidative phosphorylation and TCA cycle were higher abundant in cluster 2 tumors.

**Figure 5.**
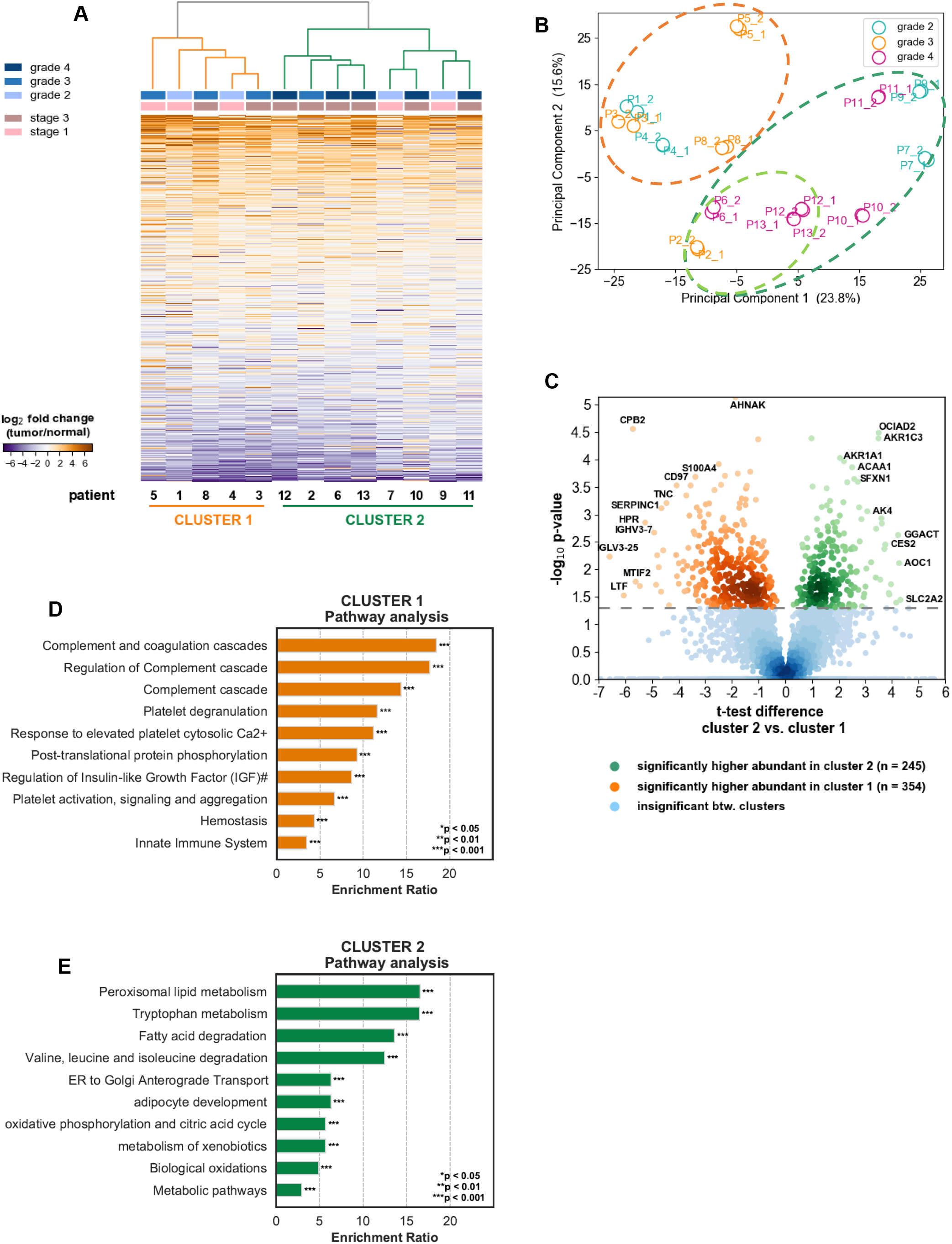
Proteome-based stratification of ccRCC tumors. **(A)** Hierarchical clustering of RCC tumors into two main clusters by expression of quantified proteins (5,799). Grade and stage information of tumors is given in column bars. **(B)** Principal Component Analysis of technical replicates of ccRCC samples. Biological variance reflects observed cluster profiles. **(C)** Volcano plot representing proteins with significantly different expression between clusters. T-test *p*-values < 0.05 were considered statistically significant. **(D-E)** Pathway analyses of cluster 1 **(D)** and cluster 2 **(E)** proteins, respectively, based on KEGG&Reactome&Hallmark50 annotations. Top 10 enriched annotations are represented. Full annotation of term marked with #: Regulation of Insulin-like Growth Factor (IGF) transport and uptake by insulin-like Growth Factor Binding Proteins (IGFBPs).

### Cluster signature is conserved in independent ccRCC mRNA and protein expression data

For the validation of the proposed clustering profile, two independent larger datasets were used. First, the TCGA-KIRC RNASeq dataset (38), consisting of 539 tumors (normalized to 72 normal tissue counts), was utilized. In general, more than 5,500 mRNA-protein pairs displayed a significant positive correlation (0.38-0.42, *p*-value <0.0001) similar to previous observations in proteotranscriptomics analysis of ccRCC (7, 22) (Figure S3B), warranting the suitability of this transcriptomics data for validating our proteome derived clustering signature. Furthermore, 565 cluster-significant proteins had an mRNA counterpart, which also demonstrated significant positive correlation (*p*-value <0.0001), with a Spearman correlation of 0.6 and a Pearson correlation of 0.42, respectively (Figure 6A). Surprisingly, proteome-transcriptome regulation discrepancy was higher for cluster 1-specific pairs (Spearman 0.49, Pearson 0.38) compared to cluster 2-specific pairs (Spearman 0.74, Pearson 0.69). This may suggest that metabolic processes in cluster 2-tumors are less prone to discordant regulation between the transcriptome and proteome level than immune and angiogenesis related processes in cluster 1-tumors.

**Figure 6.**
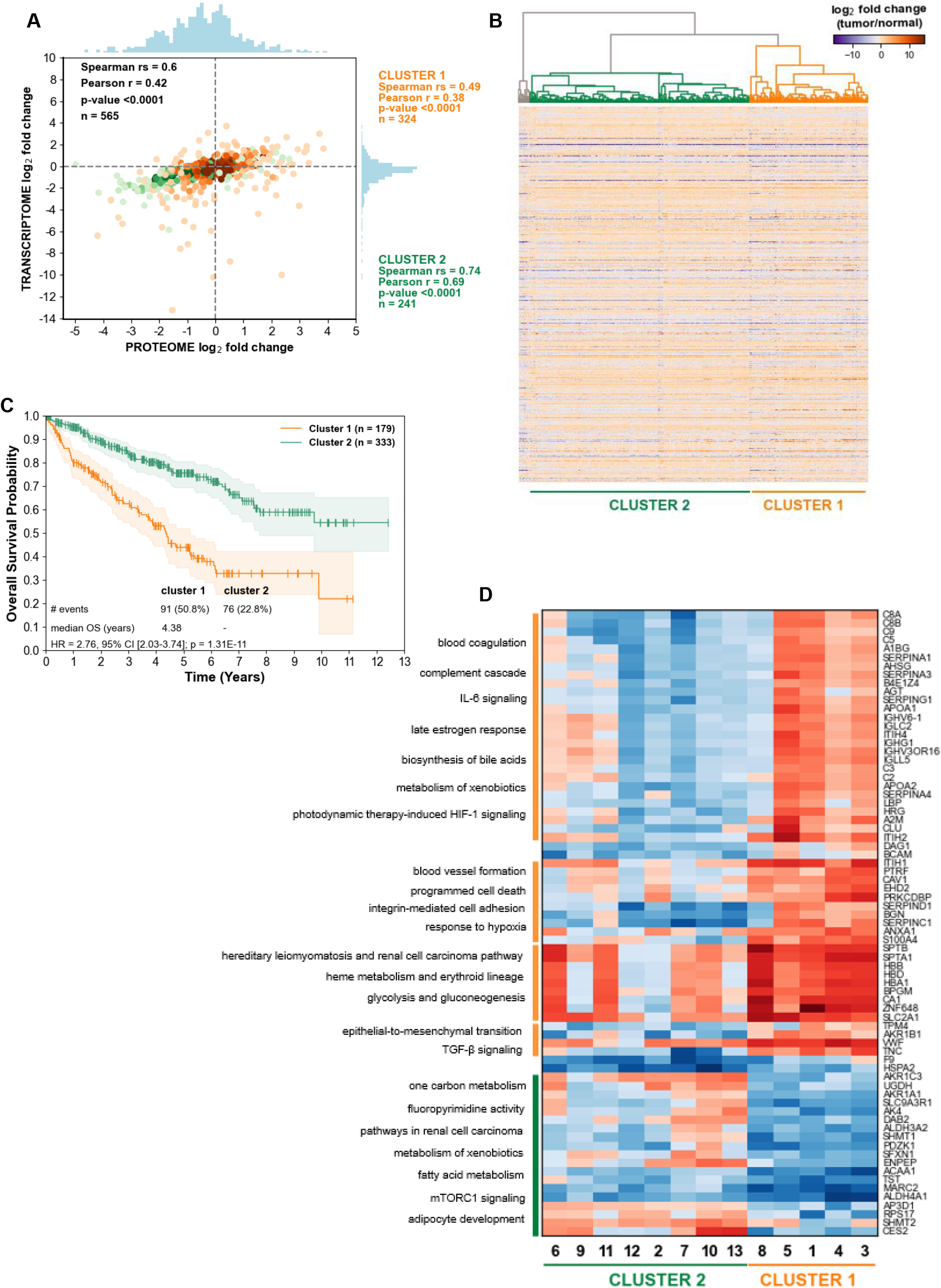
Validation of clustering signature of ccRCC tumors. **(A)** Proteotranscriptomics analysis of 565 cluster-significant proteomics-TCGA pairs comparing median log2 tumor/normal ratios. **(B)** Hierarchical clustering of TCGA-KIRC samples (n=539) by cluster-significant genes. Clustering indicates two main clusters and one outlier cluster (n=19), which was excluded from validation. Clusters were assigned based on overlap with proteome-based clusters. **(C)** Kaplan-Meier analysis for overall survival data of TCGA patients. Log-rank test indicates significant survival discrepancy between patient clusters. **(D)** Heatmap representing 73 most discriminative clustering marker genes (*t*-test difference cut-off ±2, -log_10_ *p*-value >2). Pathway analyses based on Reactome&Wikipathway_cancer annotations.

Unsupervised hierarchical clustering of the TCGA data based on the expression of these 565 mRNA/protein pairs classified the tumors into 3 clusters (“TCGA-clusters”), with 2 main clusters and 1 outlier cluster (n=19), which was excluded from the validation (Figure 6B). For cluster assignment, the significantly differentially expressed genes between the TCGA-clusters were compared with those from the proteome-clusters of this study and assigned based on their overlap. TCGA-cluster 1 was assigned with more than 84% concordance with proteome-cluster1, and TCGA-cluster 2 was assigned with more than 62% overlap with proteome-cluster2 (Figure S3C).

We further validated the clustering signature using the Clinical Proteomic Tumor Analysis Consortium (CPTAC) data for ccRCC (22), which only included patients with tumor and NATs (n=80). Of the determined cluster-significant proteins, 547 were detected in the CPTAC proteome (Figure S3D), which displayed significantly high concordance in regulation (correlations >0.82, *p*-value <0.0001). Unsupervised hierarchical clustering of the CPTAC data based on the expression of these cluster-significant proteins also classified the tumors into two main clusters (“CPTAC-clusters”) (Figure S4A). The subsequent cluster assignment resulted in an overlap of approx. 99% with cluster 1 members and 94% with cluster 2 members of this study, respectively (Figure S4B).

### Cluster 1 patients have almost three times higher risk of death than cluster 2 patients

Next, we examined the prognostic difference between the clusters using the TCGA-KIRC follow-up data. Strikingly, Kaplan-Meier analysis of the overall survival (OS) revealed highly significant discrepancy in clinical outcome between the clusters (log-rank test *p*-value <0.001) (Figure 6C, Table S1). Patients of cluster 1 had an approximately 2.8 (95% Confidence Interval (CI) [2.03-3.74]) times higher risk of death (Hazard Ratio, HR) compared to patients of cluster 2 (Figure 6C). The HR value was even higher for the recurrence-free survival (RFS) of this cohort with more than 4 times higher risk of death (95% CI [2.59-6.71]) for patients of cluster 1 (Figure S4C, Table S2). Furthermore, the 5-year survival rate of the patients differed noticeably between clusters (Figure 6C). While approximately 43% (95% CI [34.4%, 51.1%]) of the patients in cluster 1 survived 5 years post-diagnosis, more than 75% (95% CI [69.6%, 80.8%]) of the patients in cluster 2 were alive.

The relatively higher mortality rate of cluster 1 patients was also supported by the evaluation of their adverse clinicopathological features (Table S3). Remarkably, cluster 1 patients demonstrated significantly higher correlation with unfavorable clinical factors such as the presence of lymph node or distant metastasis, higher stages and grades, and recurrent disease (*p*-values <0.005). Moreover, to find out whether the proposed patient clustering was among the confounding factors having a significant impact on the survival of ccRCC patients, univariate and multivariate Cox regression analyses were performed with the follow-up data of the TCGA cohort (Table S4, Table 1). Indeed, besides the age at diagnosis (*p*-value <0.01), tumor stage (*p*-value <0.005), tumor grade (*p*-value <0.05) and the presence of distant metastasis (*p*-value <0.0001) (also the presence of lymph node metastasis in univariate analysis, *p*-value <0.0001), the cluster membership was determined as a statistically significant predictor of survival in both analyses (*p*-value <0.0001). In consideration of all parameters, the assignment to cluster 1 indicated a similarly high risk of mortality as the presence of distant metastasis (HR of 2.11 and 2.18, respectively) (Table 1).

**Table 1.** Summary of multivariate Cox Regression analysis using clinical data of TCGA-KIRC cohort (n = 490). Regression model was generated using CoxPHFitter function of lifelines library. “Not reported” entries for covariates were filtered out. CI = Confidence Interval.

| Parameter | Category setting | Hazard Ratio | Lower 95% CI | Upper 95% CI | p-value |
| --- | --- | --- | --- | --- | --- |
| Diagnosis Age | >60 vs <60 | 1.605 | 1.146 | 2.250 | 0.006 |
| Prior Cancer Diagnosis Occurrence | Yes vs No | 1.040 | 0.657 | 1.648 | 0.866 |
| Disease Stage | III+IV vs I+II | 1.888 | 1.242 | 2.869 | 0.003 |
| Histologic Grade | G3+G4 vs G1+G2 | 1.493 | 1.005 | 2.216 | 0.047 |
| Gender | Male vs Female | 0.792 | 0.565 | 1.112 | 0.179 |
| Cluster | 1 vs 2 | 2.113 | 1.500 | 2.977 | 1.88E-05 |
| Fraction Genome Altered | >50% vs <50% | 0.963 | 0.468 | 1.982 | 0.919 |
| Race Category | Asian vs Black vs White | 0.881 | 0.528 | 1.468 | 0.626 |
| Cancer Metastasis Stage Code | M1 vs M0+MX | 2.177 | 1.482 | 3.196 | 7.22E-05 |
| Lymph Node Stage | N1 vs N0+NX | 1.478 | 0.750 | 2.914 | 0.259 |

To obtain a predictive biomarker panel sufficient for cluster discrimination, we further reduced the 599 cluster-significant proteins to a subset of representative proteins by applying cut-off values on the statistics (-log_10_ *p*-value >2, *t*-test difference ±2). The resulting 73 discriminative proteins, hereafter denoted as “marker genes”, comprised of 54 cluster 1-specific and 19 cluster 2-specific proteins, respectively (Figure 6D). These marker genes were sufficient to recapitulate the significant difference in survival between the patient clusters (Figure S4D). Besides their expected role in angiogenesis and immune response related processes (Figure 5D), marker genes of cluster 1 were also associated with TGF-β related metastasis, apoptosis and HIF1α-signaling (Figure 6D). This would confirm the significant progressive and malignant clinical behavior of high-risk cluster 1 tumors. Contrarily, besides their expected primary involvement in metabolic processes (Figure 5E), marker genes of cluster 2 were also associated with mTOR signaling and adipocyte development (Figure 6D), both of which have previously been described to promote RCC progression (2, 39).

### Cluster-discriminative marker genes are druggable

To elucidate the cross-talk between marker genes of both clusters, we created a protein-protein interaction network complemented with Steiner nodes (Figure 7A, markers from cluster 1 are labelled yellow, markers from cluster 2 are labelled green). We observed that the most prominent hubs CAV1, F2R, FLOT2, CFTR, APOA1, CANX, GJA1, LGALS3, ELANE and FN1 were highly interconnected (Figure 7A, nodes with thick border). Considering that two of the hubs and most of their interactors were cluster 1-specific, we hypothesized that this underlying interconnectivity might be an attractive target for drug treatment of cluster 1 tumors. This prompted us to annotate the network with FDA-approved drugs with inhibitor function. A total of 13 out of the 87 cluster-discriminative marker genes and Steiner nodes were targets for 39 FDA-approved drugs (Figure 7B). The majority of the pharmaceutical options that are used in RCC treatment target the VEGF(R) (sunitinib, pazopanib, sorafenib, axitinib) or mTOR (everolimus, temsirolimus) pathways (22). Although these annotated drugs for the marker genes are generally applied for diseases other than cancer such as hypertension, cardiovascular conditions, pulmonary embolism, anxiety disorders, osteoarthritis or inflammatory conditions, their repurposed application in combination with the FDA-approved drugs for ccRCC might be worthwhile as a future treatment perspective. For example, two marker gene hubs are targetable with drugs, namely ELANE with alpha-1-proteinase inhibitor and CTFR with ibuprofen and dexibuprofen. Also, as shown in our study, immune system related processes play a major role in the highly lethal cluster 1 tumors. Considering that five druggable marker genes (CA1, C5, MME, ELANE and ACAA1) were involved in the innate immune system response (Figure 7A), stressing the importance of developing novel immunotherapy options is crucial for future treatment options.

**Figure 7.**
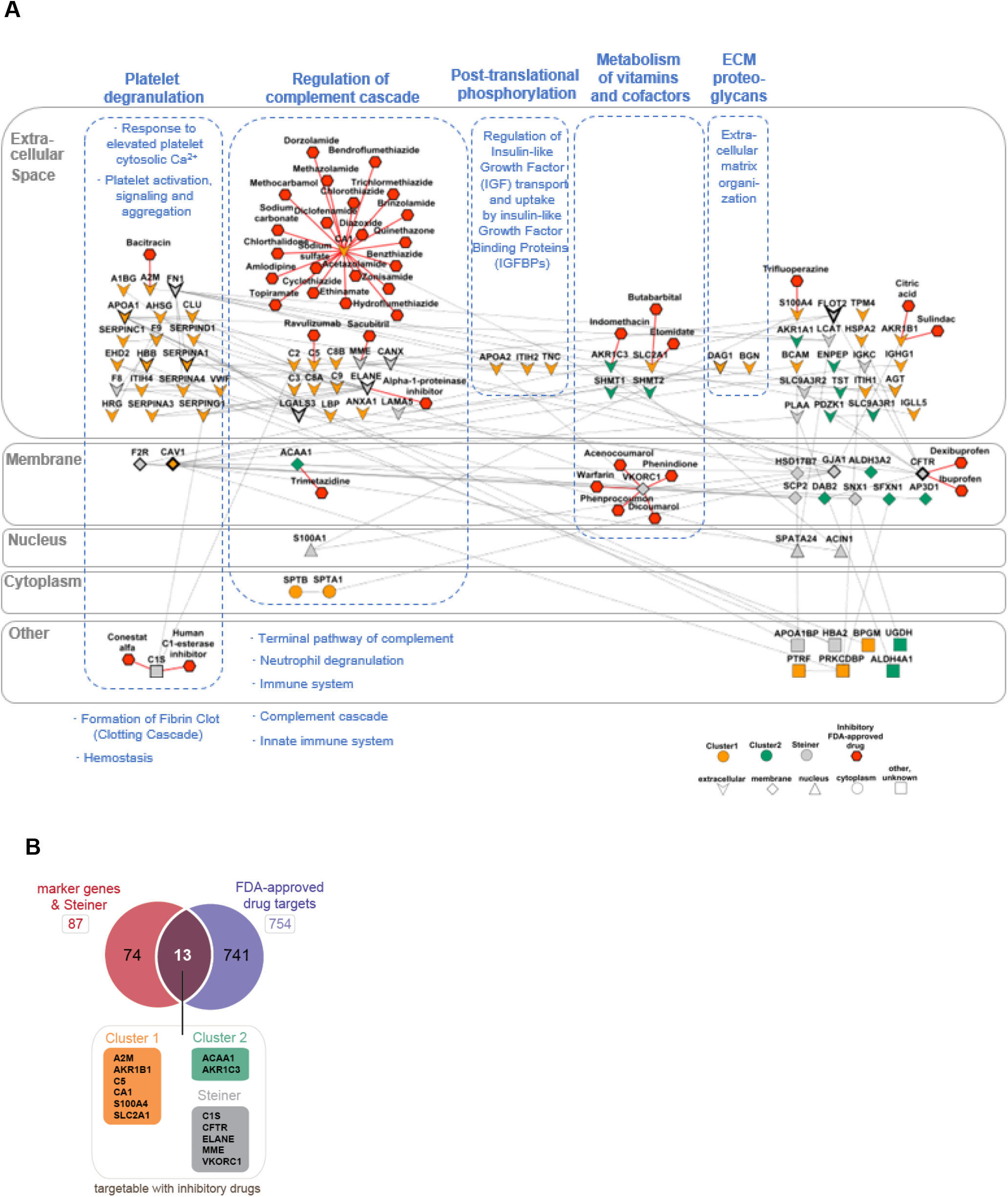
Interaction network of cluster-discriminative marker genes. **(A)** Reconstructed protein-protein interaction network with addition of Steiner nodes by using Omics Integrator. FDA-approved inhibitory drugs are linked to 13 drug targets. GO Cellular Compartment and associated Reactome pathways of proteins were retrieved by QuickGO and ClueGO, respectively. **(B)** Venn diagram showing FDA-approved drug targets among the marker genes.

## Discussion

In this study, we performed quantitative proteomics analysis of frozen ccRCC tissues and NATs and identified a total of 10,160 proteins, of which 5,799 were quantified by dimethylation (Figure 2A). Moreover, 955 of these proteins were determined as statistically significant between tumor and NATs. A remarkable undertaking for the comprehensive proteogenomic characterization of ccRCC was performed by the Clinical Proteomic Tumor Analysis Consortium (CPTAC), representing the deepest proteome coverage of ccRCC with a total of 11,355 proteins identified and 7,026 proteins quantified from 103 patients (22). Comparing both proteome datasets revealed high conformity, with 4,540 of the quantified proteins of this study (84.5%) being shared with the CPTAC study (Figure S5A) and displaying significant positive correlation (*p*-value<0.05) as indicated by a Spearman and Pearson correlation of 0.68 for each (Figure S5B). A total of 382 proteins of the significantly regulated proteins identified in the CPTAC study, were also discovered in this study, validating our data (Figure S5C). In addition, we identified 573 unique, differentially expressed proteins (Supplementary File 2), promising a distinct proteome characterization ccRCC compared to the CPTAC approach.

Out of the detected 409 significantly up-regulated proteins, we selected 4 biomarkers, namely PLOD2, FERMT3, SPARC and SIRPα that are potentially secreted, and confirmed their significantly elevated expression in ccRCC tumor tissues compared to NAT samples (Figure 3). None of the biomarkers were significantly abundant in the proposed tumor clusters (Figure 5C), which supports that these proteins are unaffected not only by intratumor heterogeneity (Figure 3E), but also by interpatient tumor heterogeneity, supporting their eligibility as universal biomarkers of ccRCC. To the best of our knowledge, SIRPα and SPARC haven’t been detected in RCC by proteomics approaches. Although PLOD2 and FERMT3 are in the list of significantly up-regulated proteins in other ccRCC proteomics studies (9, 22, 40, 41), both have not been validated by other approaches orthogonal to proteomics.

PLOD2 (Lysyl hydroxylase 2, LH2) modifies telopeptidyl lysine residues of pro-collagen α chains on the endoplasmic reticulum and is generally regulated by HIF1α, TGF-β and in RCC also by the tumor-suppressive microRNAs miR-26a-5p and miR-26b-5p (42–44). Aside from PLOD2, our PRM analysis also showed a higher abundance for the other known PLODs, PLOD1 (LH1) and PLOD3 (LH3) in ccRCC tumors (Figure 3C,D). Despite its previously reported possible secretion in lung cancer (44), we could not detect PLOD2 in urine samples (Figure 4B,C), however, it is worthwhile to investigate its suitability in the clinical setting by using a larger cohort and by measuring the urinary level of its collagen degradation products.

FERMT3, also known as Kindlin-3 and URP2, functions in hemostasis and thrombosis and plays a pivotal role in integrin-mediated cell-to-cell crosstalk as well as cell-matrix junctions (45, 46). Although it’s role in cancer remains unclear, it has been reported to be highly expressed in different types of lymphoma and breast cancer, leading to increased tumor growth, angiogenesis and metastasis (45, 46). Interestingly, FERMT3 has been reported to act as a tumor suppressor in RCC, breast tumors and melanoma based on mRNA data (45). The ambiguous role of FERMT3 in cancer warrants the need for further investigations.

SIRPα is a transmembrane protein which interacts with the ubiquitously expressed transmembrane protein CD47 (“SIRPα-CD47 axis”) (47). This cross-talk contributes to the escape of tumor cells from the immune system response (47). Our data confirms that SIRPα is located to the membrane in ccRCC tumor cells (Figure 4A), however, we could also detect SIRPα in urine samples of ccRCC patients as well as control patients, implicating that SIRPα is also a secreted protein. Given its role in immune evasion and its high abundance in ccRCC tissues, the SIRPα-CD47 axis can be considered as another promising target for immunotherapy besides nivolumab against PD-1 or ipilimumab against CTLA-4 (48).

SPARC (Secreted Protein Acidic and Rich in Cysteine) is a multi-faceted matricellular glycoprotein that is involved in tissue remodeling, morphogenesis, bone mineralization (49). While it functions as a tumor suppressor in neuroblastomas, colorectal and ovarian cancer (49, 50), it has been reported to be elevated on the mRNA level in sarcomatoid RCC, as well as most of ccRCC tumors (70%) (51). SPARC has been reported to be expressed only by tumor-associated stromal cells such as vascular endothelial cells and fibroblasts, but not by cancer cells themselves (51). In line with this, we observed for SPARC an intense staining in the intratumor stroma (Figure 4A). Furthermore, we detected a significantly elevated abundance of SPARC in urine samples of ccRCC patients making it a promising tool for clinical assays (Figure 4B,C).

Here, we also propose a biomarker panel for the appropriate classification of ccRCC tumors based on the underlying molecular phenotypes, since classification solely based on histological or morphological features of tumors and subsequent treatment decisions can lead to varying treatment response (20, 52). In this study, we identified two ccRCC tumor clusters that differed in their molecular features. Our clustering signature is distinct from previously suggested ones such as the widely investigated transcriptomics based ClearCode34 grouping system, which has only 5 proteins in common (Figure S5D) (21). Both the ClearCode34 as well as our classification system showed similar hazard ratios for the overall survival between the patient clusters (HR 2.4 and 2.2, respectively). However, regarding the recurrence-free survival, our clustering signature showed a 4.2 fold difference in the risk of death between the clusters while the difference was 2.3 fold between the ClearCode34 clusters. Furthermore, our clustering signature distinguishes from the proteomics based classification system ccRCC1-3 (22), which showed malignant features intermixed across the 3 identified clusters, while in this study we showed that highly malignant processes such as epithelial-to-mesenchymal transition (EMT), HIF1α signaling, innate immune response and angiogenesis were condensed in cluster 1 tumors accounting for the relatively worse survival outcome (Figure 6C,D), while cluster 2 tumors had a low-risk profile and a more favorable prognosis.

Advanced RCC is currently treated with monotherapies or combination therapies of immune checkpoint inhibitors and VEGF or mTOR inhibitors in first- and second-line strategies (22, 53). Hypothesizing from our data, immunotherapy as first-line treatment could possibly be more effective for the highly immunogenic cluster 1 tumors (Supplementary File 3), while mTOR inhibitors could possibly augment treatment response for cluster 2 tumors (Supplementary File 3).

Furthermore, we identified 13 cluster-discriminative marker genes targetable by repurposed FDA-approved drugs (Figure 7A) that can provide alternative treatment strategies to meet the need for novel anticancer therapeutics (54). One highly elevated EMT marker in cluster 1 tumors is the signaling protein S100A4, which is a target of the antipsychotic drug trifluoperazine (Figure 7A). It has been shown that trifluoperazine can be repurposed as an anticancer drug and successfully prevent or minimize metastatic spread in different cancers by inhibiting S100A4 through protein oligomerization (55, 56). Targeting S100A4 could also be advantageous for reducing hypoxic signaling in cluster 1 tumors, since it was revealed as HIF1α target (Supplementary File 3). Another gene highly overexpressed in cluster 1 tumors upon hypoxic signaling is SLC2A1 (Supplementary File 3), also known as glucose transporter 1 (GLUT 1). This marker gene can be targeted by the intravenous anesthetic etomidate (Figure 7A), which has been reported to reduce GLUT1-mediated glucose transport and associated tumor growth in lung cancer (57, 58). SLC2A1 has also been revealed to be related to angiogenic processes such as coagulation and platelet degranulation, as well as the marker CA1 (Supplementary File 3), which participates in hypoxia-induced glucose transport, angiogenesis and pH regulation (59). Multiple inhibitory FDA-approved sulfonamides such as acetazolamide, methazolamide, dichlorophenamide, dorzolamide and brinzolamide can target CA1 (Figure 7A), and have been reported to counteract metastatic dissemination in different RCC cell lines (60).

Taken together, our comprehensive proteomics analysis identified 4 novel biomarkers for the detection of ccRCC. Future directions should entail the investigation of the role of these biomarkers in ccRCC tumorigenesis and metastatic spread, as well as the application of SPARC as an analytical tool in clinical assays. We further uncovered 2 distinct molecular phenotypes of ccRCC tumors and identified a cluster-discriminative biomarker panel targetable by several repurposed FDA-approved drugs. Future directions warrant the investigation of tumor-oriented intervention by combining these novel therapeutics with current medical treatment options of RCC.

## Supporting information

Patient info

List of significantly regulated proteins in tumors

List of enriched pathways in patient clusters

## Acknowledgements

We gratefully acknowledge the KUPAM and OMICS facilities of Koç University. We thank Ozgur Kurt and Gulsum Citak from Koç University School of Medicine for their essential help in pathological assessment of tissue samples. We further thank Dr. Serkan Kir for valuable scientific comments on the results. The results shown here are in whole or part based upon data generated by the TCGA Research Network: https://www.cancer.gov/tcga. Data used in this publication were generated by the Clinical Proteomic Tumor Analysis Consortium (NCI/NIH).

## Author Contributions

A.S. and N.O. designed the research plan. O.A., M.C.K. and S.E. accrued patients and supervised patient sample collection. A.S. performed proteomics experiments, statistical analysis and data analysis. A.T.S. performed immunoblotting experiments (with contributions from A.S.), and related data analysis. A.A. performed pathological assessment of tissues, conducted IHC experiments and wrote the IHC protocol. S.B. performed pathological assessment of tissues. O.A., M.C.K., S.E., A.A. and T.E. consulted on clinical questions. N.T. supervised bioinformatics analysis. N.O. conceived and supervised the study. A.S. wrote the manuscript with inputs from N.O. All authors discussed the results and commented on the manuscript.

