## Supplementary figures for "Quantitative proteomics identifies secreted diagnostic biomarkers as well as tumor-dependent prognostic targets for clear cell Renal Cell Carcinoma"

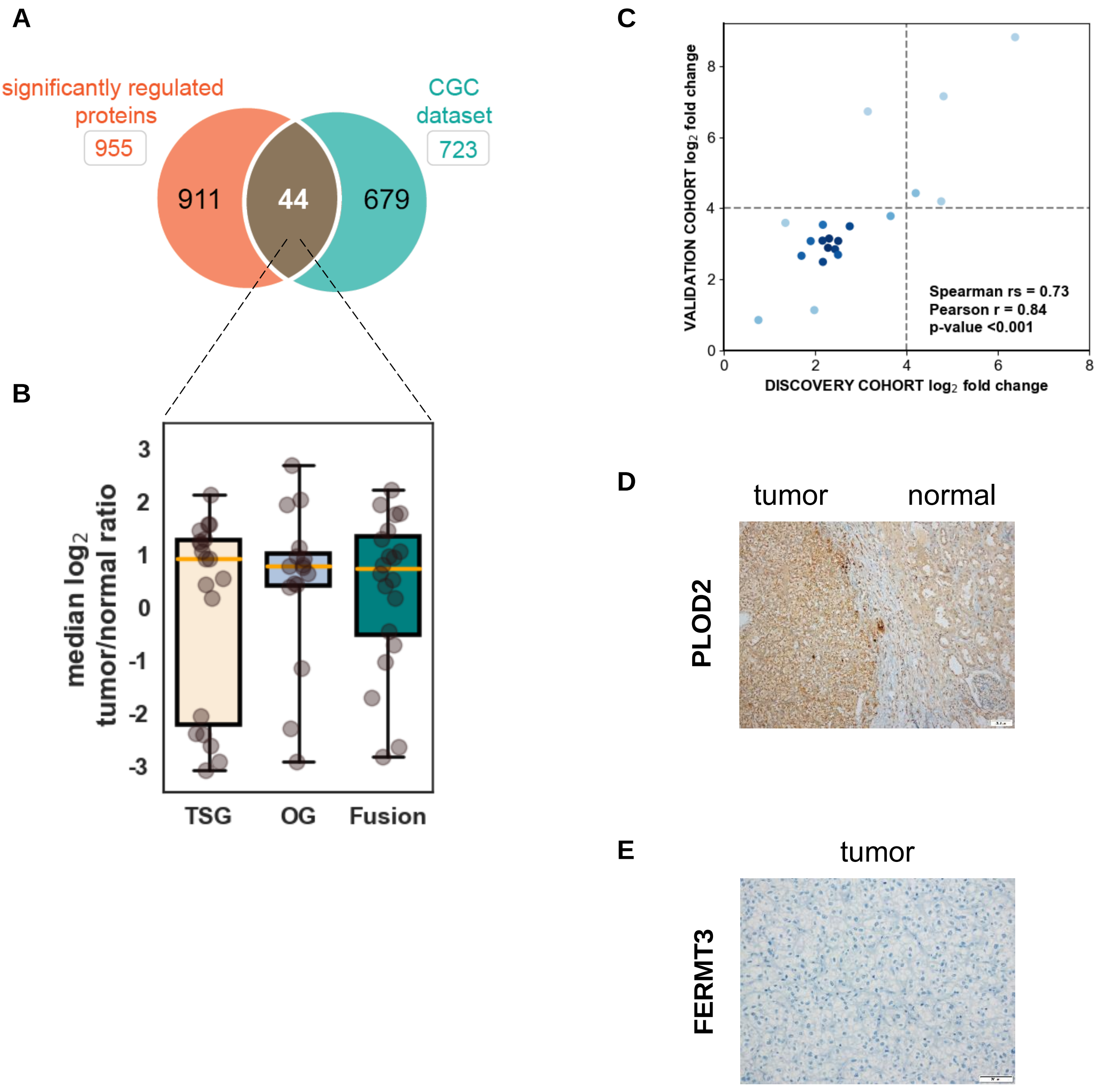


**Figure S1. Characteristics of significantly regulated proteins.** (**A**) Venn diagram depicting number of cancer genes among the significantly regulated proteins in ccRCC tumors retrieved from the COSMIC Cancer Gene Census database. (**B**) Distribution of significantly regulated cancer genes categorized by their function. Cancer genes might be allocated to multiple functional categories. (**C**) Correlation of PRM tumor/normal ratios between discovery and validation cohort. (**D**) IHC staining of PLOD2 expression in transitional region of tumor and normal tissue. (**E**) IHC staining of FERMT3 expression in tumor tissue.

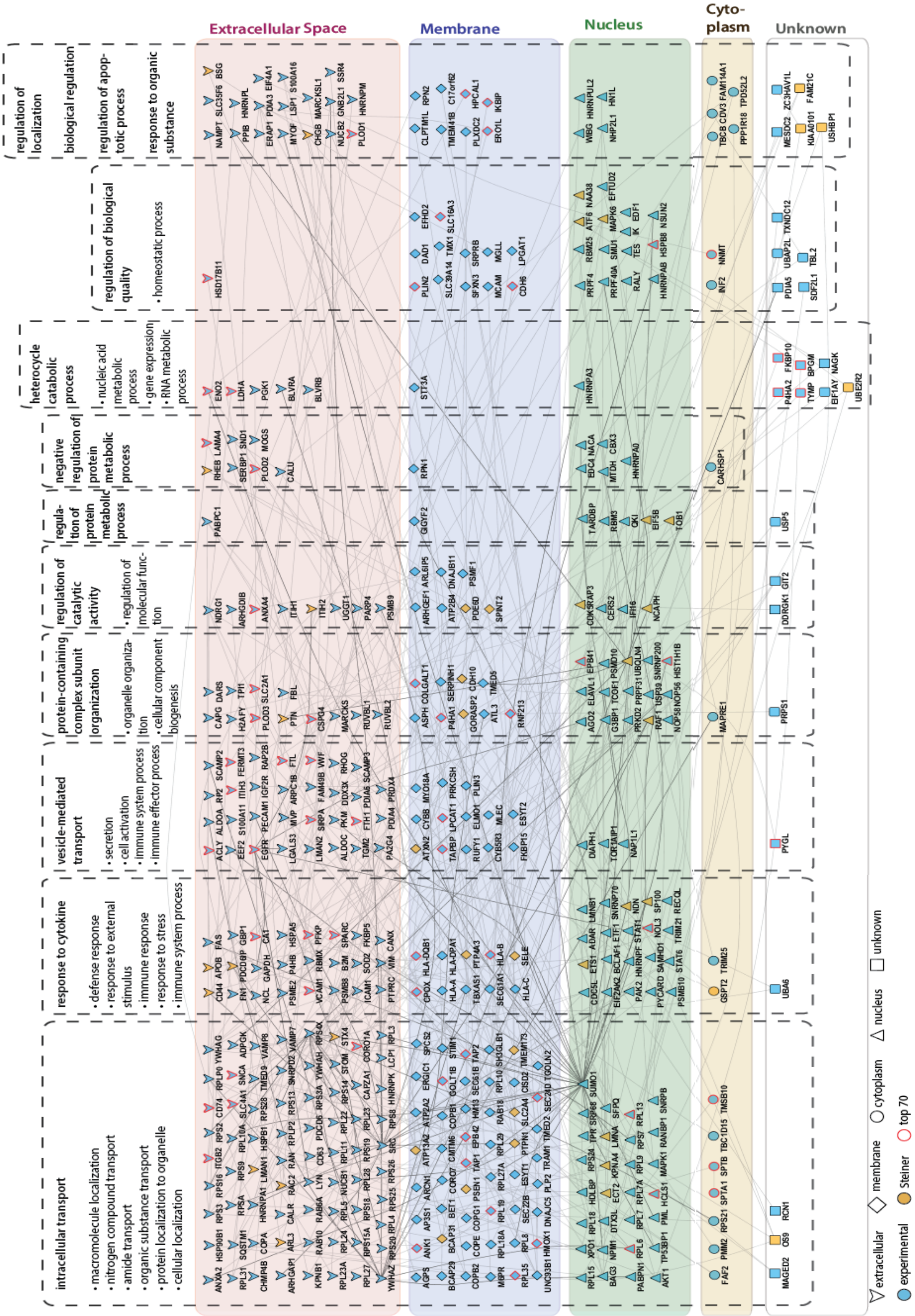


**Figure S2. Complete protein-protein interaction network of significantly up-regulated proteins in ccRCC tumors.** Edge annotations including intermediate Steiner nodes created using Omics Integrator. Associated GO Cellular Compartment and Biological Process information of proteins were retrieved by QuickGO and ClueGO, respectively.

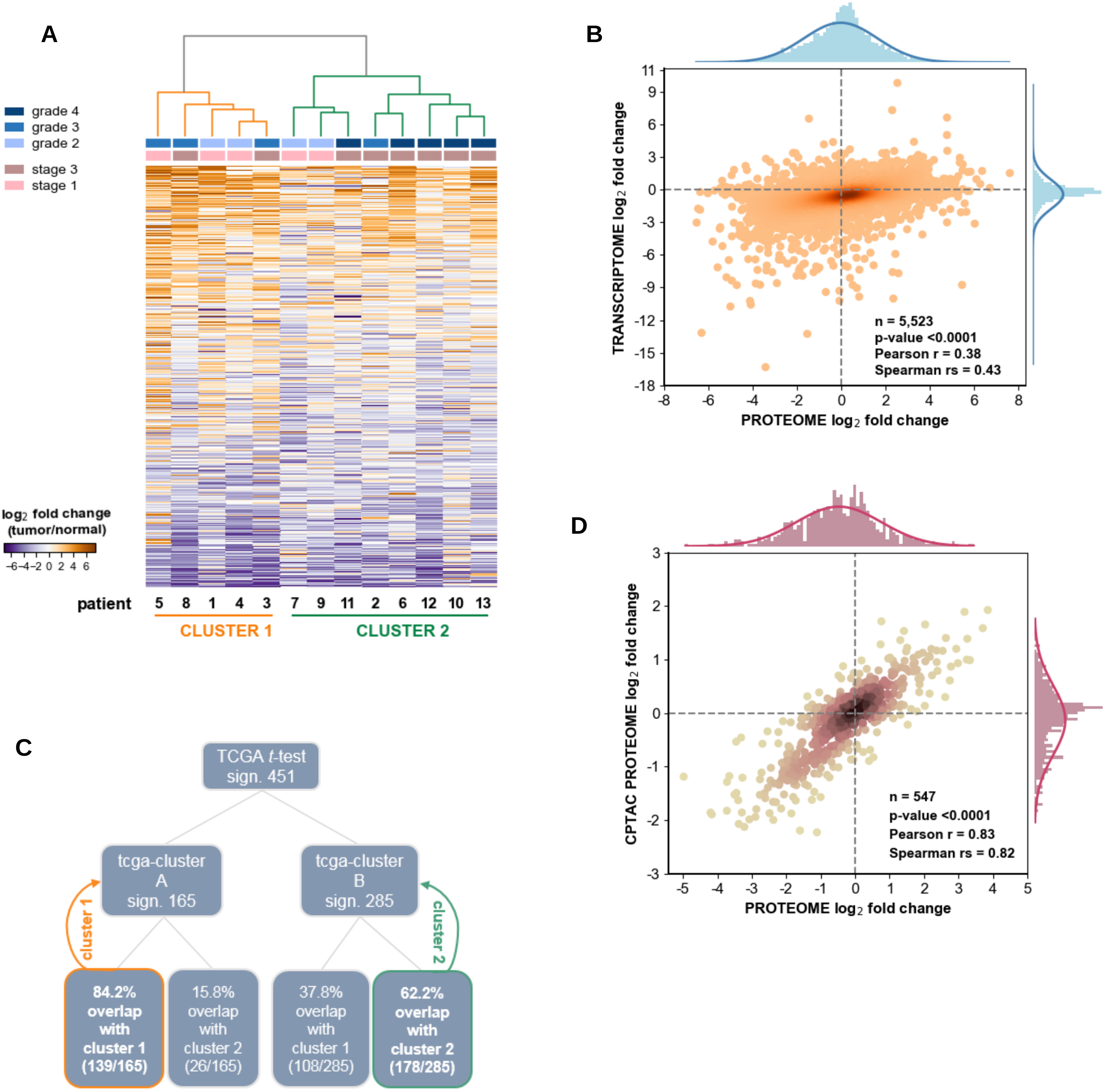


**Figure S3. Robustness and validation of clustering signature of ccRCC tumors.** (**A**) Hierarchical clustering of tumors into expected two main clusters using expression data of 866 high-variant proteins. (**B**) Proteotranscriptomics analysis of total proteome and total TCGA-KIRC data by median $\log_2$ tumor/normal ratios. (**C**) Cluster assignment of TCGA clusters based on overlap with proteome clusters. (**D**) Correlation analysis between proteome data and CPTAC data for 547 cluster-significant proteins based on median $\log_2$ tumor/normal ratios.

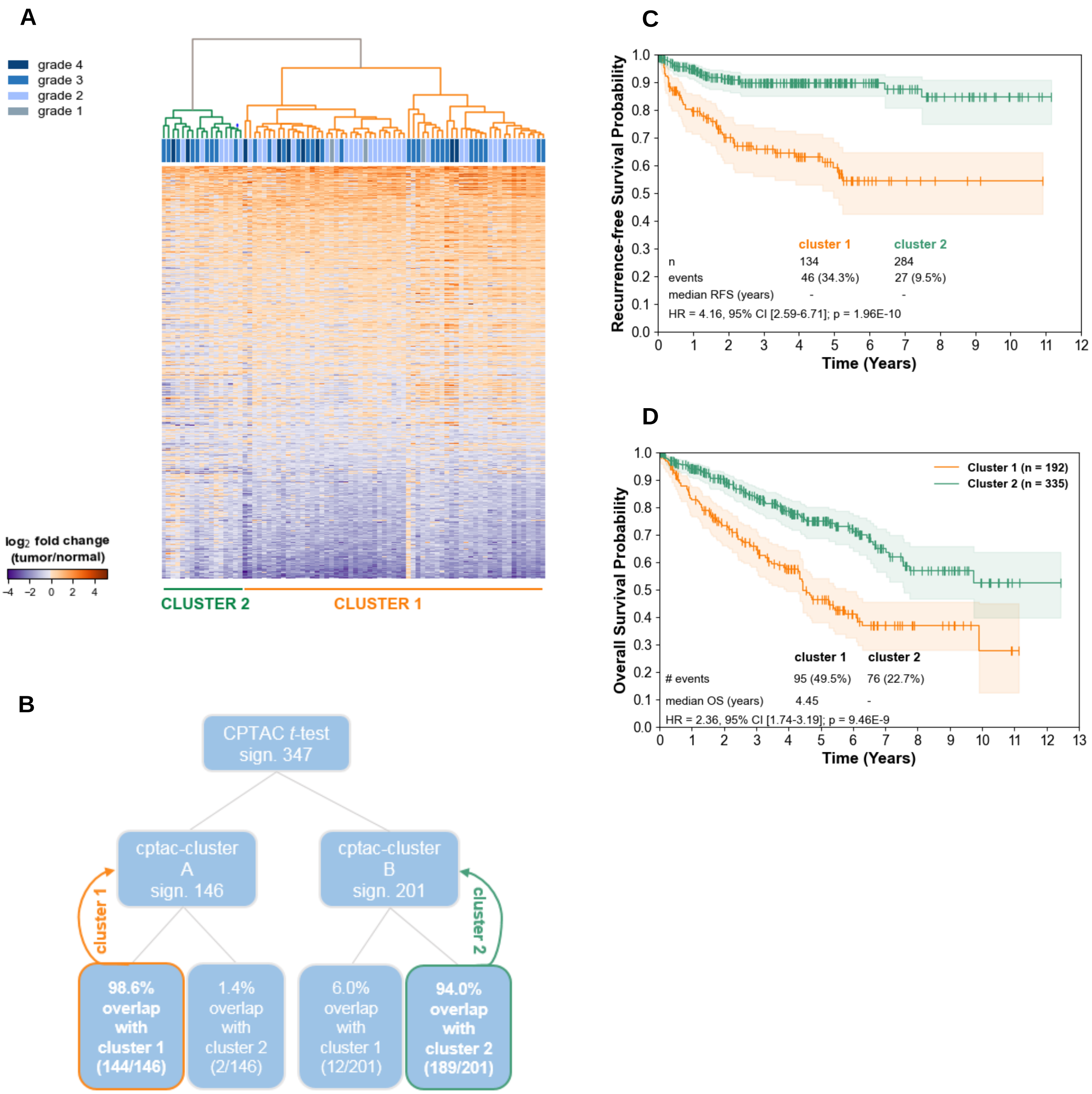


**Figure S4. Validation and survival analysis of clustering signature.** (**A**) Hierarchical clustering of CPTAC samples (n=80) by cluster-significant proteins (n=547) into two clusters. (**B**) Cluster assignment of CPTAC clusters based on overlap with proteome clusters. (**C**) Kaplan-Meier analysis and log-rank test of recurrence-free survival data of TCGA patients indicate significant survival discrepancy between patient clusters. (**D**) Kaplan-Meier analysis and log-rank test of overall survival data of TCGA patients based on clustering by 73 cluster-discriminative marker genes which are sufficient to recapitulate significant survival discrepancy between patient clusters.

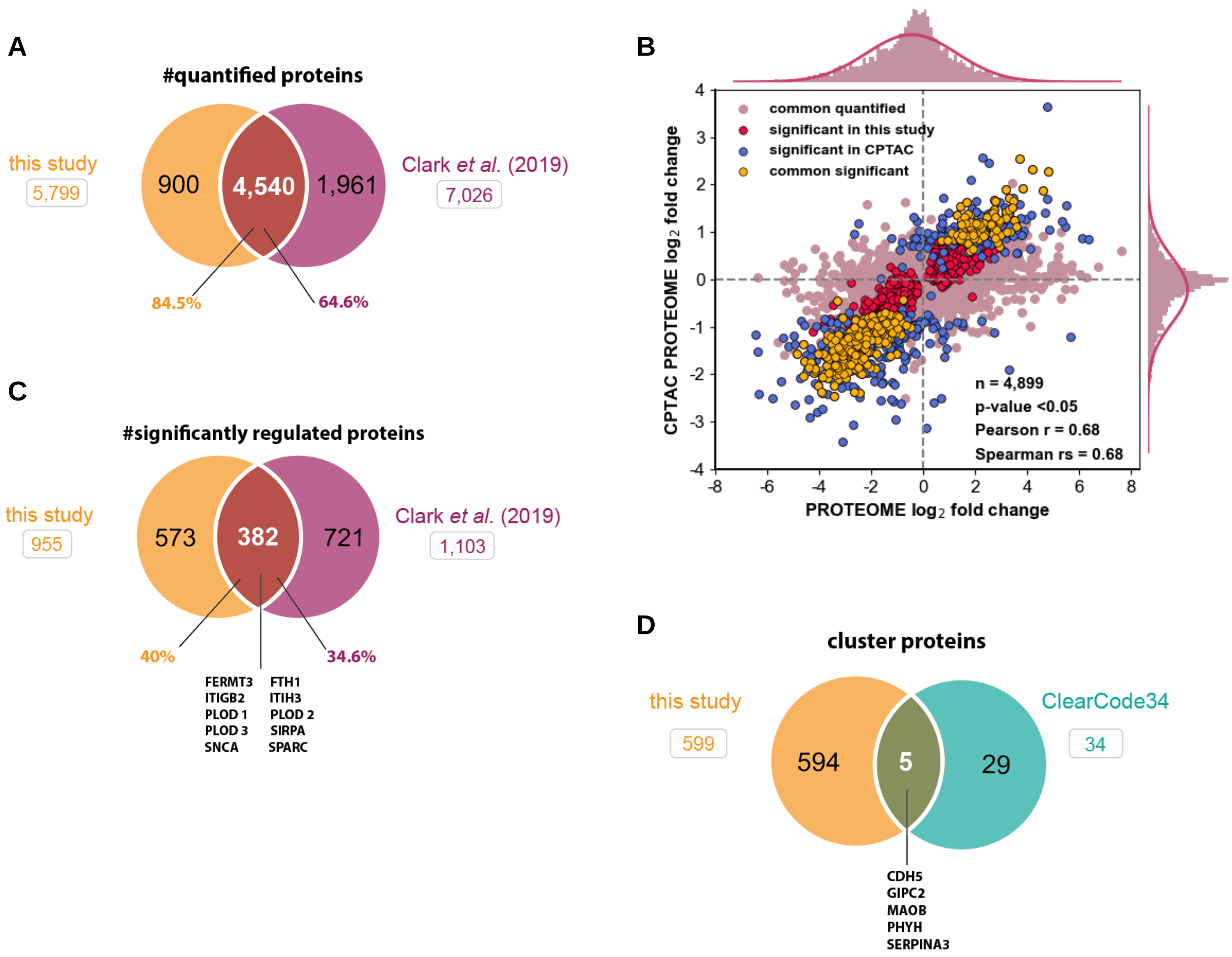


**Figure S5. Comparison of proteome data with CPTAC study and ClearCode34 classification system.** (**A**) Venn diagram showing number of common quantified proteins with CPTAC dataset. (**B**) Correlation between proteome data and CPTAC data for all quantified proteins based on median $\log_2$ tumor/normal ratios. (**C**) Venn diagram showing number of common significantly regulated proteins with CPTAC dataset. (**D**) Venn diagram showing number of common cluster-significant proteins with ClearCode34 predictive dataset.
