## Supplementary tables for "Quantitative proteomics identifies secreted diagnostic biomarkers as well as tumor-dependent prognostic targets for clear cell Renal Cell Carcinoma"

**Table S1. Summary of Overall Survival (OS) for ccRCC patient clusters using TCGA-KIRC cohort (n = 520).** Python library lifelines used for Kaplan Meier survival curve generation. 95% confidence intervals shown in parentheses.

| | no. of patients | no. of death events | percentage of death events | median survival (years) | 5-year survival rate | 10-year survival rate | max. follow-up time (years) |
|---|---|---|---|---|---|---|---|
| **Cluster 1** | 182 | 93 | 51.1% | 4.38 [3.48-5.24] | 42.9% [34.4%-51.1%] | 21.5% [6.9%-41.2%] | 11.14 |
| **Cluster 2** | 338 | 76 | 22.5% | - | 75.7% [69.6%-80.8%] | 54.7% [42.4%-65.5%] | 12.43 |

**Table S2. Summary of Recurrence-Free Survival (RFS) for ccRCC patient clusters using TCGA-KIRC cohort (n = 418).** Python library lifelines used for Kaplan Meier survival curve generation. 95% confidence intervals shown in parentheses.

| | no. of patients | no. of death events | percentage of death events | median survival (years) | 5-year survival rate | 10-year survival rate | max. follow-up time (years) |
|---|---|---|---|---|---|---|---|
| **Cluster 1** | 134 | 46 | 34.3% | - | 59.1% [48.3%-68.4%] | 54.4% [42.5%-64.8%] | 10.92 |
| **Cluster 2** | 284 | 27 | 9.5% | - | 89.7% [85.1%-92.9%] | 84.7% [74.9%-90.9%] | 11.15 |

**Table S3. Comparison of clusters of TCGA-KIRC patients by clinicopathological parameters (n = 534).** For diagnosis age the ttest_ind function, for the other parameters the fisher_exact function of the scipy.stats library was used.

| | Cluster 1 | Cluster 2 | Fold frequency difference cluster 1 vs cluster 2 | *p*-values |
|---|---|---|---|---|
| **Size** | 181 | 333 | - | - |
| **Mean diagnosis age** | 61.5 | 60.0 | - | 0.181 |
| **Metastasis stage M1** | 24.3% (44) | 9.9% (33) | 2.45 | 2.63E-05 |
| **Lymph node stage N1** | 6.7% (12) | 1.2% (4) | 5.58 | 2.02E-03 |
| **Disease Stage I + II** | 44.2% (80) | 68.5% (228) | 0.64 | 1.13E-07 |
| **Disease Stage III + IV** | 55.2% (100) | 30.9% (103) | 1.79 | 1.03E-07 |
| **Histologic Grade 1 + 2** | 29.3% (53) | 54.7% (182) | 0.54 | 3.7E-08 |
| **Histologic Grade 3 + 4** | 69.6% (126) | 44.7% (149) | 1.56 | 6.7E-08 |
| **Disease free** | 39.8% (72) | 66.9% (223) | 0.59 | 3.26E-09 |
| **Recurred/Progressed** | 35.4% (64) | 18.3% (61) | 1.93 | 2.49E-05 |
| **Fraction genome altered > 50%** | 6.1% (11) | 2.4% (8) | 2.54 | 0.048 |

**Table S4. Summary of univariate Cox Regression analysis based on Overall Survival using clinical data of TCGA-KIRC cohort (n = 534).** Regression model was generated with the CoxPHFitter function of lifelines library. “Not reported” entries for covariates were filtered out. CI = Confidence Interval.

| Parameter | Category setting | Hazard Ratio | Lower 95% CI | Upper 95% CI | *p*-value | Cohort size |
|---|---|---|---|---|---|---|
| **Diagnosis Age** | >60 vs <60 | 1.857 | 1.351 | 2.551 | 1.36E-04 | 531 |
| **Prior Cancer Diagnosis Occurrence** | Yes vs No | 0.834 | 0.542 | 1.285 | 0.411 | 531 |
| **Disease Stage** | III+IV vs I+II | 3.863 | 2.810 | 5.311 | 8.69E-17 | 528 |
| **Histologic Grade** | G3+G4 vs G1+G2 | 2.700 | 1.911 | 3.816 | 1.81E-08 | 523 |
| **Gender** | Male vs Female | 0.939 | 0.688 | 1.281 | 0.691 | 531 |
| **Cluster** | 1 vs 2 | 2.756 | 2.029 | 3.741 | 8.37E-11 | 512 |
| **Fraction Genome Altered** | >50% vs <50% | 1.030 | 0.526 | 2.017 | 0.931 | 523 |
| **Race Category** | Asian vs Black vs White | 0.813 | 0.484 | 1.362 | 0.431 | 524 |
| **Cancer Metastasis Stage Code** | M1 vs M0+MX | 4.483 | 3.287 | 6.114 | 2.6E-21 | 529 |
| **Lymph Node Stage** | N1 vs N0+NX | 3.909 | 2.116 | 7.223 | 1.35E-05 | 531 |
